# Using herbarium genomics to understand the history of a global plant invasion

**DOI:** 10.64898/2026.03.31.715310

**Authors:** Ramona-Elena Irimia, Cosimo Posth, Ella Reiter, Tatiana Tondini, Marc W. Schmid, Uta Grünert, Yaolin Guo, Takahide Kurosawa, Michihito Ohta, Juraj Paule, Ann Bogaerts, Dmitar Lakušić, Ana Ržaničanin, Alessia Guggisberg, Christina L. Richards, Oliver Bossdorf

## Abstract

Invasive plants are dynamic eco-evolutionary systems characterized by rapid spread and evolutionary change. Herbarium genomics offers a powerful way to study these processes across broad spatial and temporal scales. We generated low-coverage shotgun-sequencing data from 152 herbarium specimens of the invasive Japanese knotweed species complex (*Reynoutria*), collected across its native range in Japan and China, and its introduced ranges in Europe and North America, spanning 200 years of global spread. Introduced populations of *R. japonica*, *R. japonica var. compacta* and *R. sachalinensis* showed reduced genetic diversity compared to native populations and were genetically most similar to Japanese specimens, indicating Japan as the source of all three introductions. European and North American *R. japonica* were genetically highly similar and closely related to local *R.* × *bohemica* hybrids, suggesting that these hybrids originated post-introduction, through hybridization and subsequent introgression from the *R. japonica* parent. A few *R.* × *bohemica* hybrids in the UK and France shared a multilocus lineage with *R. sachalinensis*, indicating an independent but ecologically less successful hybridization event involving introgression from *R. sachalinensis*. A genetically distinct *R.* × *bohemica* from Japan was closely related to a Japanese *R. sachalinensis* specimen, suggesting that hybridization also occurs in the native range. Introduced *R*. *japonica* has remained genetically uniform for ∼200 years reflecting long-term founder effects. The dominance of *a* single *R. japonica* lineage supports the ‘general-purpose genotype’ hypothesis and the importance of vegetative reproduction in its spread. Herbarium genomics thus uncovers the origin and global spread of Japanese knotweed, providing direct molecular evidence of long-term plant invasions.

## Introduction

The repeated human-mediated introductions of species to geographical locations outside of their native ranges offer replicated opportunities for understanding how demographic and evolutionary processes shape spatial genetic diversity and population differentiation in colonizing species. This can help to inform about species’ ability to adapt to new habitats and predict their evolutionary dynamics and invasive success.

The genetic diversity and structure of introduced populations are influenced by several genetic processes during and after introduction, including multiple introductions and admixture of different sources, inter-specific hybridization and genetic drift (Welles & Dlugosch 2018; Hodgins et al., 2025). In species with mixed reproductive systems, the prevailing mode of reproduction together with the associated dispersal mechanism, are often the primary determinants of genetic structure. These processes can have both positive and negative effects on local genetic variation, with consequences for the fate of colonizers (Welles & Dlugosch 2018). Systematic comparisons of genetic diversity in native versus invasive populations showed that the loss of genetic diversity in introduced populations is less frequent and less intense than previously expected (Dlugosch & Parker 2008), and that the so called “genetic paradox of invasions” (Estoup et al., 2016) may have been overstated. The potential for adaptative evolution to drive invasion seems to depend more on the type of genetic variation than its overall amount. For example, populations with low diversity in neutral markers can still maintain high genetic variation for traits that are ecologically important. Additionally, the genetic architecture underlying these traits plays a crucial role in shaping adaptive evolutionary responses (Dlugosch et al., 2015). Plant traits and performance can also be profoundly impacted by changes in genome dynamics in response to novel environments. Genomic redundancy resulting from whole-genome duplication (polyploidy) or divergent expression of duplicated genes (homeologs) can create functional plasticity even in the absence of genetic variation among individuals (Yoo et al., 2014; Nieto Feliner et al., 2020). Such structural and functional changes of genomes along with a variety of non-genetic effects can affect heritable variation and hence response to novel environmental conditions (reviewed in Mounger et al., 2021). Understanding when this variation fuels evolutionary change during invasion requires identifying sources of genetic variation, tracing invasion routes and testing different evolutionary and ecological hypotheses (Estoup & Guillemaud 2010; Barker et al., 2017).

Introduction histories of invasive species are typically inferred based on historical occurrence records or indirect methods based on patterns in population genetics data (Estoup & Guillemaud 2010; Dormontt et al., 2014; Nierbauer et al., 2016; Vallejo-Marín et al., 2021; Bieker et al., 2022). However, such genetic inferences are usually derived from extant populations that have been shaped by complex genetic processes during introductions, which makes it challenging to reconstruct distant historical events and early stages of invasion (Dormontt et al., 2014; Martin et al., 2014). Looking back in time might be possible through natural history collections (*e.g.*, herbaria) that document invasions from introduction through different subsequent stages. Recent advances in sequencing technologies now allow the successful retrieval of genome-wide data in herbarium and museum collections and thus open the possibility to study the genetic composition of invasions through time and capture evolutionary events as they occurred (Kistler et al., 2020; Burbano & Gutaker 2023; Kim et al., 2023). Some previous studies have already provided important insights into invasion pathways (*Senecio madagascariensis*: Dormontt et al., 2014; *Macfadyena unguis-cati* and *Jatropha gossypiifolia:* Prentis et al., 2009; *Mimmulus guttatus*: Vallejo-Marín et al., 2021), rapid adaptation during range expansion and colonization (*Ambrosia artemisiifolia*: Bieker et al., 2022; Battlay et al., 2023; *Sisymbrium austriacum*: Vandepitte et al., 2014) and shift in interactions with natural enemies (Bieker et al., 2022) for some highly invasive plant species and have proven the usefulness of these methods.

Invasive knotweeds (*Reynoutria* spp., Polygonaceae) are among the 100 worst invasive species (Lowe et al., 2004). The spread of *Reynoutria japonica* (Japanese knotweed) has frequently been considered an example of the “genetic paradox of invasion”, where an introduced species is successful across a wide range of habitats despite little or no genetic diversity (Estoup et al., 2016). Across western Europe and the British Isles, populations of *R*. *japonica* appear to consist of clonal replicas of a single octoploid (8x) female plant (Hollingsworth & Bailey 2000a; Mandák et al., 2004; Holm et al., 2018; Jugieau et al., 2024; Krebs et al., 2009; Tiébré et al., 2007). However, genetic studies in central-eastern Europe (Bzdęga et al., 2012) found moderate genetic variation among *R*. *japonica* populations, challenging the idea of continent-wide genetic uniformity. The genetic structure of north-eastern US populations of *R*. *japonica* is more complex, comprising both male and female plants (Del Tredici 2017; Forman & Kesseli 2003) and different genotypes (Dodson 2009; Wymer et al., 2007; Grimsby et al., 2007; Gammon & Kesseli 2010; VanWallendael et al., 2021), which suggests multiple introductions to this area (Del Tredici 2017) and different invasion histories of the two continents. Nonetheless, studies in the Mid-Atlantic (Richards et al., 2012) and on the western coast (Gaskin et al., 2014) of the US have found only one widespread octoploid (8x) *R*. *japonica* genotype that is identical to the invasive lineage in western Europe and the British Isles (Hollingsworth & Bailey 2000a; Mandák et al., 2004; Krebs et al., 2009; Zhang et al., 2016). In contrast to *R*. *japonica*, the other taxa in this species complex, *R*. *sachalinensis* and *R.* × *bohemica* are genetically much more diverse across Europe and North America and comprise both male and female individuals as well as several unique genotypes possibly resulting from independent hybridization (Mandák et al., 2004; Tiébré et al., 2007; Krebs et al., 2009; Gaskin et al., 2014; VanWallendael et al., 2021). Compared to its wide geographic introduced range, *R*. *japonica* has a more restricted native range, limited to Japan, Eastern China, Korea and Taiwan, which may reflect control by natural enemies or plant competitors, or biogeographical barriers that constrain range expansion (Bailey 1989, Beerling et al., 1995). Surveys of Asian *R*. *japonica* have detected considerable genetic diversity among native populations (Pashley 2003; Inamura et al., 2000; Zhou et al., 2003; Zhang et al., 2024) along with high variability in ploidy levels (4x, 6x, 8x and 12x, reviewed in Mandák et al., 2003) and morphological traits (Kim and Park 2000).

The introduction history of invasive knotweeds has been previously studied in a series of herbarium-based studies that aimed to identify introduction events and map the spread and current taxa distribution in Europe (Bailey & Conolly 2000; Mandák et al., 2004; Thiébaut et al., 2020) and North America (Barney et al., 2006). However, there are so far only few molecular investigations of both native and introduced populations (Pashley 2003; Desjardins et al., 2023; Zhang et al., 2024), often with a limited sampling range (but see the recent paper by Zhang et al., 2024) and using low-resolution markers (*e.g.*, cpDNA, AFLP, GBS) without a reference genome. Consequently, the specific origins of introduced genotypes, along with the timing and frequency of propagule introduction and their effects on the dynamics of this invasion remain poorly understood. Here, we conducted shotgun sequencing on historical herbarium specimens of Japanese knotweed *s*.*l*. collected between 1828 and 2018, including the earliest known introduction sites in Europe and North America, along with putative source populations from the native range in Asia. We aimed to: (a) assess the prevalence of clonal spread vs. sexual reproduction in the invasion, (b) verify the native origin of introduced populations and determine whether the invasion resulted from multiple independent introduction events or a single event, (c) examine how genetic diversity in introduced populations compares to that of native populations, (d) explore the role of hybridization in the success of the invasion. We predicted that an increased number of genetic markers, along with the recently published reference genome for *R. japonica* (Wang et al., 2025a) and expanded temporal sequencing across a broad geographic range would provide the power to detect multiple introductions events, including unsuccessful introductions (*e*.*g*., genotypes that failed to establish) and uncover cryptic genetic diversity that may influence the invasion dynamics.

## Materials and Methods

### Study system

Japanese knotweed *sensu lato* (Polygonaceae) is a group of highly troublesome polyploid invasive species including *Reynoutria japonica* Houtt. (with the alpine ecotype *R. japonica* var. *compacta* (Hook.f.) J.P.Bailey), *Reynoutria sachalinensis* Nakai (Giant knotweed), hybrids between the two (*Reynoutria* × *bohemica* Chrtek & Chrtková, Bohemian knotweed), two horticultural cultivars (*R. japonica* ‘Variegata’ and *R*. *japonica* f. *rosea* Satomi or ‘Crimson Beauty’, Forman & Kesseli 2003) and crosses and backcrosses among the various taxa. All these taxa are gynodioecious, consisting of hermaphroditic and male-sterile (female) individuals. The hermaphrodites are self-incompatible and function poorly as female parents, so they generally act as male plants (Bailey 1989). *Reynoutria japonica* and *R*. *sachalinensis* are native to south-eastern Asia and were introduced to the temperate northern hemisphere and Australasia ca. 200 years ago as garden ornamentals (Bailey & Conolly 2000; Desjardins et al., 2023). The simultaneous introduction of two *Reynoutria* species facilitated hybridization between *R*. *japonica* and *R*. *sachalinensis* and resulted in the formation of the hybrid *R*. × *bohemica,* which is more aggressive and spreads faster than its parents (Parepa et al., 2014; Jovanović et al., 2018). The Japanese knotweed *s.l.* species complex exhibits considerable ploidy variation. In its introduced range, *R. japonica* is predominantly octoploid (8x) (Richards et al., 2008; VanWallendael et al., 2021; Desjardins et al., 2023; Irimia et al., 2025) whereas *R. japonica var. compacta* and *R. sachalinensis* are mainly tetraploid (4x), the latter occasionally occurring as hexaploid (6x) or octoploid (8x) (Mandák et al., 2003). The hybrid *R.* × *bohemica* displays four cytotypes: hexaploid (6x) —the most widespread cytotype globally (Irimia et al., 2025); tetraploid (4x) and octoploid (8x) —reported only from the UK and Czech Republic (Mandák et al., 2003); and decaploid (10x) —documented in artificial hybrids in the US and in a limited number of natural populations in the UK (Bailey 2013). In the wild, crosses can occur both within and among different ploidy levels (Bailey 2013).

### Herbarium sampling

We obtained leaf material from 152 knotweed herbarium specimens from 56 natural historical plant collections worldwide, including 97 individuals of *R*. *japonica*, nine of *R*. *japonica* var. *compacta*, 22 of *R*. *sachalinensis,* 22 of the hybrid *R*. × *bohemica* and two of the garden cultivar *R*. *japonica* f. *rosea*. The sampling was broad and covered the species’ distribution in its native range in SE Asia (50 accessions; 1828-2010) and the introduced range in Europe (52 accessions; 1850-2018) and North America (50 accessions; 1891-2009) (Fig. 1, Table S1). For the specimens that we received on loan (European collections) we sampled leaf tissue in our lab in Tübingen, Germany, whereas for herbaria outside Europe the curators carried out the sampling and sent us the material. Prior to sampling, we determined the taxonomic identity of the specimens using the key of Zika and Jacobson (2003) and stored the leaf material at room temperature in individual paper bags until the DNA extractions.

**Figure 1.**
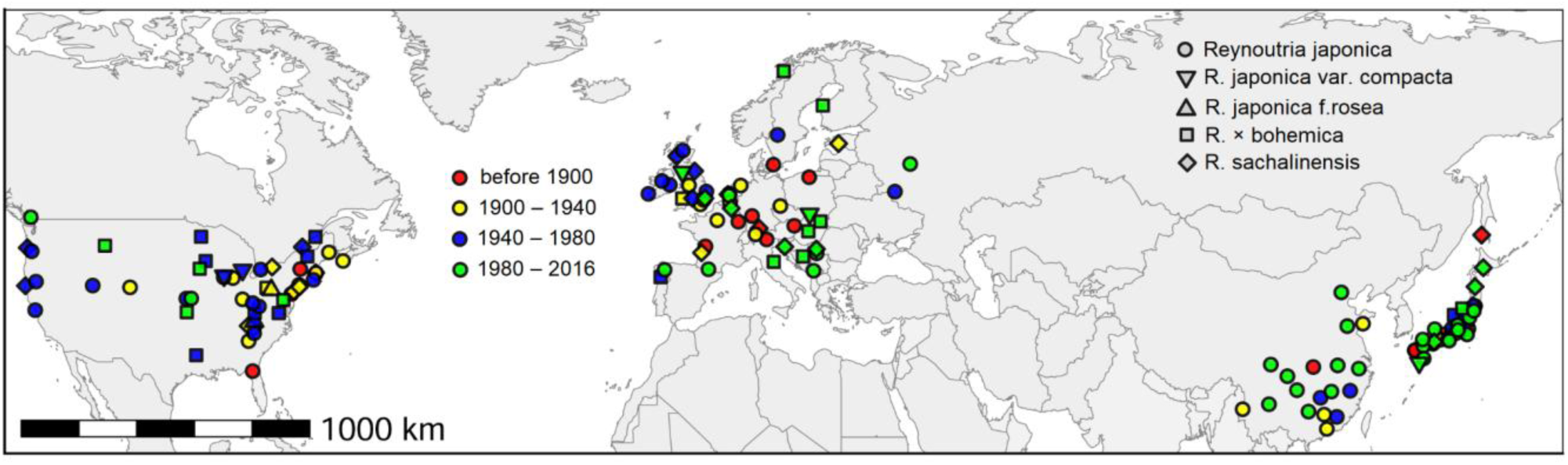
Geographic origins of the 155 samples (152 herbarium specimens + three modern samples) of Japanese knotweed *s*.*l*. included in this study. Symbols represent different taxa, while colors indicate collection periods. The species is native to Southeastern Asia (China and Japan) but has been introduced to Europe and North America where it is regarded as invasive.

### DNA isolation from herbarium specimens

We used approximately 10 mg of leaf tissue for each DNA extraction. We ground each sample into a fine powder using a benchtop mixer mill (Retsch MM 400) at a frequency of 20Hz for 1.5 min and using 3 mm stainless-steel beads for sample disruption and homogenization. This process was carried out in a lab that does not perform PCR amplification. We extracted total genomic DNA (gDNA) in a dedicated ancient DNA “clean room” facility at the Institute for Archaeological Sciences at the University of Tübingen, using DNeasy Plant Mini kits (Qiagen) and following the protocol of Lattore et al., 2020. For DNA lysis, a solution of 1.2 ml N-phenacylthiazolium bromide (PTB)-based mix and 48μl proteinase K was added to each sample. To enhance cell lysis, we incubated the samples overnight at 37°C in a lab rotator. After incubation, we centrifuged the samples and transferred them into a binding buffer containing guanidine hydrochloride which increases the recovery of ultra-short DNA molecules (<50bp, Lattore et al., 2020). After washing twice with the washing buffers supplied by the DNeasy Plant Mini kit, we eluted the DNA in EB (elution buffer) in a final volume of 50μl and stored it at −20°C until further processing. We processed a maximum of 10 samples in parallel, including one negative control without any plant tissue. We checked the DNA integrity on a 4150 TapeStation System (Agilent Technologies) using the D1000 Screen Tape and Reagents kit and determined the gDNA concentration of each sample with a Qubit 3.0 Fluorometer (Life Technologies) and the Qubit^TM^ 1x dsDNA HS assay kit (Invitrogen).

### Double stranded (ds) DNA library preparation, quantification and sequencing

We built double stranded DNA libraries following the protocol by Meyer and Kircher 2010. Libraries were prepared without uracil DNA glycosylase treatment (ds_nonUDG: 20 samples) for screening of historical DNA or with partial uracil DNA glycosylase treatment (ds_halfUDG: 132 samples, among these 11 samples were processed in triplicate as technical replicates, Table S1) to reduce but not eliminate the amount of deamination at the end of historical DNA fragments. We used a starting DNA template of 15-20 μl (*e.g.,* 200 ng DNA) and built library batches of 20 or 22 samples each, including a negative control without template DNA. All steps for library preparation were done in the ancient DNA clean room facility mentioned before. Briefly, we used a mixture of DNA polymerases (T4 Polynucleotide Kinase and T4 Polymerase, New England BioLabs) to conduct blunt-end repair of the DNA fragments followed by reaction purification with MinElute columns (Qiagen), Solexa adapter ligation: (Read 1 sequencing primer: 5’-ACACTCTTTCCCTACACGACGCTCTTCCGATCT-3’ and Read 2 sequencing primer: 5’-GTGACTGGAGTTCAGACGTGTGCTCTTCCGATCT-3’), using Quick Ligation^TM^ kit (New England BioLabs) and adapter fill-in using Bst DNA polymerase (New England BioLabs). For quality control, we included a library preparation negative control (no DNA template) in two of the library batches. We quantified the amplifiable library molecules by qPCR using the IS7 (sequence: ACACTCTTTCCCTACACGACGC) and IS8 (sequence: GTGACTGGAGTTCAGACGTGTGC) primers with a DyNAmo SYBR Green qPCR kit (Thermo Fisher Scientific) on a LightCycler 480 (Roche). We then double-indexed (8bp long barcodes, Table S2) each DNA library in 1-14 parallel reactions using PfuTurbo DNA polymerase (Agilent). After setting the indexing PCR reaction in the clean room, we moved to the modern lab for library amplification. We indexed the libraries using an initial denaturation at 95°C for 2 min and 10 cycles of 95°C for 30 sec, 58°C for 30 sec, and 72°C for 1 min, followed by a final elongation at 72°C for 10 min. We pooled the indexed products for each library and purified them with MinElute columns (Qiagen), followed by elution in 50μl TET (1M Tris-HCl, 0.5M EDTA, 0.05% Tween-20, pH 8.0) and quantification by qPCR using the IS5 (sequence: AATGATACGGCGACCACCGA) and IS6 (sequence: CAAGCAGAAGACGGCATACGA) primers. We checked the fragment distribution and library concentration of the indexed libraries on the TapeStation and then diluted the individual DNA libraries to 10nM and prepared an equimolar pool of all libraries for shotgun sequencing. The libraries, including two negative library controls, were sequenced on an Illumina NovaSeq platform at Genewiz (Azenta Life Sciences, Leipzig, Germany) using a 2×150bp pair-end configuration kit.

### Raw data processing & genotyping

We generated 547.7 GB of whole genome sequencing raw data (WGS) and ∼4.6B paired end reads. As a first step in data processing, we used the EAGER pipeline (Peltzer et al., 2016) to map, authenticate and assess the quality of DNA in our samples. We ran the pipeline using default parameters, with the following steps: quality control of sequencing (FastQC analysis), removal of sequencing adapters and merging of paired-end data (Adapter Removal), mapping reads to the reference genome (Wang et al., 2025a) using BWA with a seed length of 16500, setting the amount of allowed mismatches in the alignment to 0.01 and a mapping quality filter of 30, removal of PCR duplicates (DeDup), calculation of ancient DNA C-to-T damage (MapDamage2.0), estimation of library complexity and generation of summary statistics. Across all samples, the proportion of endogenous plant DNA accounted for 70–84% of all reads, and the C-to-T damage at the first DNA bases was approximately 0.05%-0.06% for the non-UDG treated libraries and about 0.03% for the half-UDG treated libraries.

To improve regional coverage in China, we co-analyzed three modern samples from that area (collected in 2023) with the 152 herbarium samples. These three accessions had previously been generated using a WGS approach (Illumina NovaSeq 6000), and we downloaded them from GenBank (SRR25018594, SRR25018595, SRR25018596, see Table S1). For SNP calling, we quality-trimmed the reads with fastp (version 0.20.0, Chen et al., 2016) and aligned them to the *R. japonica* reference genome using only one representative for each of the two main sub-genomes, chromosome set A & E (National Genomics Data Center: PRJCA021875; *R*. *japonica* plant accession from Wuhan, Hubei Province, China, Wang et al., 2025a), using BOWTIE2 (Langmead and Salzberg 2012; version 2.4.5). We removed duplicates with Picard tools (version 2.18.25) and extracted alignment and duplication statistics from the console output or logs. We identified SNPs with bcftools (version 1.18, Danecek et al., 2021) by evaluating genotype likelihoods from aligned reads and calling variants at positions with statistical support for an alternative allele relative to the reference. Specifically, we filtered variant calls for a minimum Phred quality score of 20 and kept only biallelic loci with a coverage of at least 10 in 50% of all individuals, which resulted in an original data set of 2,409 SNPs (2,382 nuclear and 27 plastid).

### Multi-locus lineage identification

We used the original dataset (i.e., 2,409 SNPs) to determine which samples shared the same multi-locus lineages (MLL). Specifically, we used the genetic distances observed between technical replicates (e.g., within-sample distances, see Table S1) as a background distribution for random noise. We calculated pairwise genetic distances as the fraction of alleles that differed between two samples (ignoring SNPs for which one or both samples did not have data available). We considered samples with genetic distances smaller than the average distance between technical replicates plus three times the standard deviation to belong to the same multi-locus lineage (we chose the cut off for differences that define identity to a given MLL as 0.0576 for all taxa). We used these distances as edges in a network to identify distinct MLLs using a modularization algorithm (Blondel et al., 2008). Samples/replicates from the same individual were merged by majority call. To visualize MLL groups and overall genetic structure of different taxa, we conducted a principal component analysis using the PCA function in the *adegenet* 1.3-1 package (Jombart & Ahmed 2011) in R version 4.5.1 (R Core Team, 2025).

### Genetic divergence

We excluded SNPs that exhibited inconsistencies across technical replicates (i.e., differences within replicate pairs) to filter out variants with potential errors, further reducing our dataset to 1,964 SNPs (1,937 nuclear and 27 plastid). We computed pairwise Nei’s genetic distances (hereafter abbreviated as *D*) between taxa using the *dist.genpop()* function in the *adegenet* package and visualized the resulting distance matrix as a heatmap with *ggplot2* (Wickham 2016).

### Genetic diversity and regional allele pools

To characterize the genetic diversity of each taxon in each geographic region, we calculated multiple components of genomic diversity across the 1,964 error filtered SNPs. We estimated observed heterozygosity (H_o_), mean nucleotide diversity (π) and the fixation coefficient (F_IS_) using the *hierfstat* 0.5-11 package (Goudet & Jombart 2024) in R. For the two parental species, *R. japonica* and *R. sachalinensis*, we also calculated the total, shared, and private allele counts across geographic regions using functions from the *adegenet* 1.3-1 package (Jombart & Ahmed 2011). We then constructed a Jaccard similarity matrix to quantify the similarity of allele pools between pairs of regions and visualized the distribution of shared and unique alleles across regions using a Venn diagram generated with *ggvenn* (Yan 2025). To account for unequal sample sizes among taxa, we estimated genetic diversity using a rarefaction approach. For each taxon with at least five individuals, we randomly subsampled five individuals without replacement. We repeated this procedure 200 times, calculated the diversity statistics for each replicate and reported the final values with means and the confidence intervals based on the rarefaction replicates.

### Phylogenetic relationships

Finally, we used the error filtered 1,964 SNPs across 150 individuals (Fig S1) to infer evolutionary relationships across individuals and taxa. We conducted all phylogenetic analyses using the maximum-likelihood (ML) approach method implemented in PhyML 3.0 (Guindon et al., 2010, http://www.atgc-montpellier.fr/phyml/). Specifically, we employed the GTR + R (Free Rate model: accounts for sites evolving at different speeds) + F (uses observed nucleotide frequencies) nucleotide substitution model and the SPR (Subtree Pruning and Regrafting) algorithm with a parsimony-based filter to explore tree topologies, starting from a BioNJ (Gascuel 1997) or a minimum parsimony (MP) tree. We assessed branch support using a non-parametric Shimodaira-Hasegawa-like (SH) approximate likelihood ratio test (Kishino & Hasegawa 1989). As input, we converted the SNP data into an aligned FASTA file in PHYLIP format where each column of the SNP table was concatenated to form a sequence. We visualized the phylogenetic tree using iTOL (Letunic and Bork 2024, https://itol.embl.de/). To better understand maternal lineage patterns, we repeated the phylogenetic analyses using a subset of 27 plastid derived SNPs (see Fig. S1 and accompanying supplementary text).

## Results

### Multi-locus lineage identification

We identified six distinct multi-locus lineages (MLLs) (Fig. 2A, Table S1) among the 153 analyzed samples. MLL 1 included all but one of the Chinese *R*. *japonica* individuals, together with several *R*. *japonica* accessions from central (Nagano and Aichi prefectures) and southern Japan (Mie, Oita, Nara, Kochi, Tokushima and Tottori prefectures) as well as two *R*. *japonica* f. *rosea* individuals from North America. MLL 2 consisted of four *R*. *japonica* individuals from the southernmost part of Japan (Oita, Miyazaki, Kagoshima and Yamaguchi prefectures). MLL 3, the largest lineage, comprised all *R*. *japonica* individuals from Europe and North America, as well as the majority of *R*. × *bohemica*. Additionally, it included nine *R*. *japonica* individuals from Japan (collected across central Japan, with one from Kyushu Island near Nagasaki) and one individual from China (Shandong province). MLL 4 comprised all *R*. *sachalinensis* individuals but one, along with three *R*. × *bohemica* accessions (two from UK and one from France). MLL 5, the smallest, included a *R*. × *bohemica* accession from the west coast of Honshu Island and a *R*. *sachalinensis* accession from Shikoku Island in southern Japan. Finally, MLL 6 included the *R*. *japonica* var. *compacta* group, with specimens from Mt. Fuji and surrounding prefectures, as well as from Europe and North America. Additionally, one accession from Japan (Tokyo: 1985, FKSE7311), identified on the herbarium label as *R*. *japonica* var. *hachidyoensis*, displayed a unique genotype that could not be assigned to any of the MLL groups and as a result was excluded from the MLL PCA analysis but included in the taxa PCA (Fig. 2B, Table S1).

**Figure 2.**
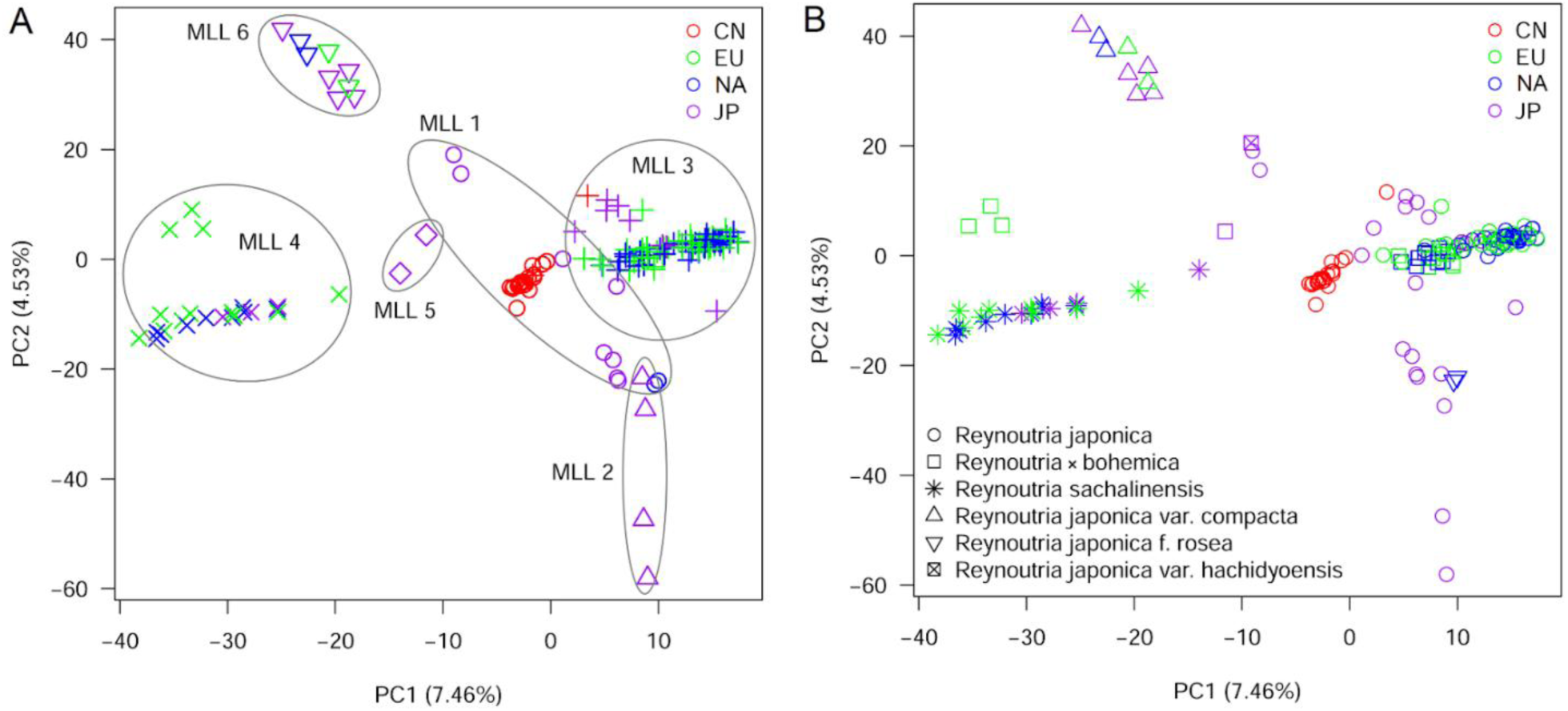
Principal component analysis (PCA) plot showing the assignment of the 153 individuals of Japanese knotweed *s*.*l*. (150 herbarium specimens and three modern samples from China) based on 1,964 SNPs to A) different multi-locus lineages (MLL) and B) different *Reynoutria* taxa. Each point in the PCA is an individual. Geographic regions are abbreviated as follows: CN - China, EU - Europe, NA - North America (US and Canada) and JP - Japan.

### Genetic divergence

When we quantified the genetic distances among taxa within and between geographic regions, we found very low regional genetic differentiation for *R. japonica*, *R. japonica var. compacta* and *R. sachalinensis* (Fig. 2B, Fig. 3). Specifically, *R. japonica* individuals from Europe and North America were nearly identical and genetically much closer to some Japanese samples than to Chinese samples, supporting the hypothesis of a single introduction event of Japanese origin, whereas Chinese *R. japonica* individuals were more differentiated from the other regions. There were similarly close relationships for *R*. *sachalinensis* as well as *R. japonica* var. *compacta* individuals from different regions, also indicating single introductions from Japan. The situation was more complex in the *R.* × *bohemica* hybrids, where hybrids from continental (non-UK) Europe and North America again exhibited virtually no genetic differentiation, suggesting a common origin. However, both hybrids from Japan and the UK were more differentiated from the EU/NA lineage, and from each other, consistent with further, independent hybridization events in the native range, and within the introduced European range.

**Figure 3.**
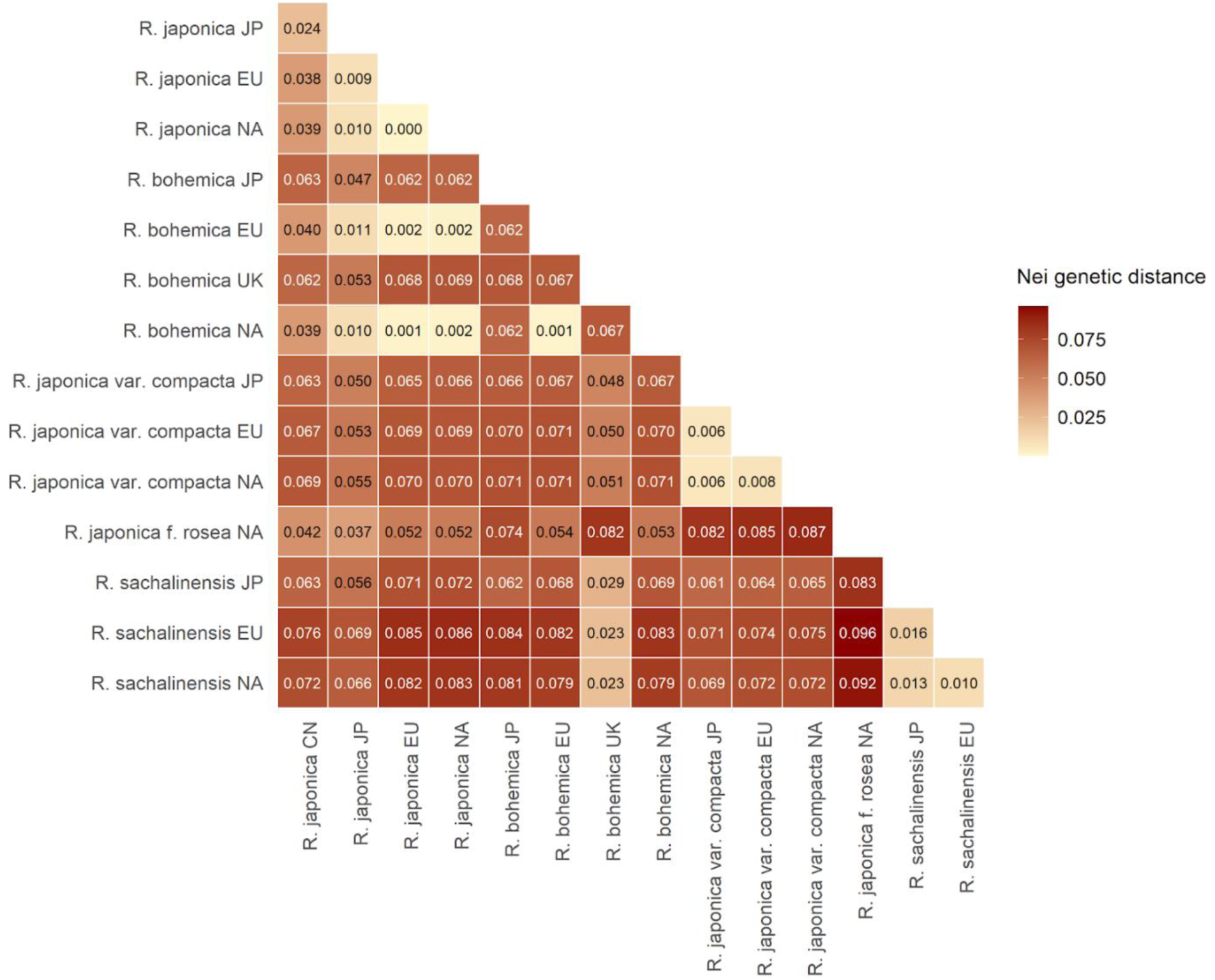
Genetic distances between different taxa in the Japanese knotweed *s*.*l*. complex, within and across different geographic regions (CN = China, EU - Europe, JP = Japan, UK = United Kingdom, NA = North America). The values in the cells are Nei’s genetic distances. The Japanese group of *R*. *sachalinenis* includes one specimen from Russia, and the UK group of *R*. × *bohemica* one from France.

When we examined genetic distances between different taxa, the most striking patterns were the different relationships of *R.* × *bohemica* hybrids from Japan vs. UK vs. other introduced regions with their putative parental taxa. The hybrids from continental Europe and North America were genetically similar to *R. japonica* from these regions and from Japan, while the hybrids from the UK were much closer to *R. sachalinensis* than the EU/NA hybrids (Fig. 2B, Fig. 3), indicating different scenarios of introgression and backcrossing in these different hybrid lineages. The hybrids from Japan, in contrast, were equally distant to both parental species. All geographic origins of *R. japonica* var. *compacta* were substantially differentiated from *R. japonica* and *R. sachalinensis*, but with slightly lower genetic distance to *R. japonica* from Japan. Finally, the garden cultivar *R. japonica* f. *rosea* was also more closely related to Japanese *R*. *japonica* than to other taxa.

### Genetic diversity

Native *R. japonica* from China and Japan showed the highest genetic diversity. In comparison, nucleotide diversity was 63% and 50% lower in Europe, and 65% and 52% lower in North America than in China and Japan, respectively (Table 1). The situation was similar in *R. sachalinensis* where European and North American plants had 36% and 34% lower nucleotide diversity than plants from Japan. In *R. japonica var. compacta* only the US individuals were genetically less diverse than the Japanese and European ones. Overall, levels of genetic diversity were very similar between the European and North American introduced ranges, indicating similar genetic bottlenecks during introduction. The only exception was the small group of *R.* × *bohemica* individuals from the UK (see also previous section) which tended to be more genetically diverse than other hybrids from Europe or North America. The values for expected heterozygosity (H_0_) showed a smaller reduction compared to nucleotide diversity, except in *R. sachalinensis* where H_0_. increased by 13% in European and North American range compared to native Japan (Table 1). Interestingly, fixation coefficients (F_IS_) were positive only in Japanese individuals of *R. japonica* and *R. sachalinensis*, whereas they were strongly negative in introduced ranges of all other knotweed taxa (Table 1).

**Table 1.** Genetic diversity statistics (mean and 95% CI) for historical Japanese knotweed *s*.*l*. individuals (different taxa) in the native range (China, Japan) and introduced ranges in Europe and North America based on 1,964 single nucleotide polymorphism loci. For groups with five or more individuals, we used rarefaction analysis to calculate diversity metric estimates. N = number of individuals, Ho = observed heterozygosity, π - nucleotide diversity and FIS -.fixation coefficient.

| Taxa | Region | N | $H_0$ | $\pi$ | $F_{IS}$ |
| --- | --- | --- | --- | --- | --- |
| *R. japonica* | China | 21 | 0.080 [0.078 — 0.083] | 0.076 [0.075 — 0.078] | -0.025 [-0.042 — -0.007] |
|  | Japan | 22 | 0.049 [0.048 — 0.050] | 0.056 [0.055 — 0.057] | 0.114 [0.099 — 0.129] |
|  | Europe | 30 | 0.044 [0.043 — 0.044] | 0.028 [0.028 — 0.029] | -0.543 [-0.548 — -0.538] |
|  | North America | 25 | 0.043 [0.043 — 0.043] | 0.027 [0.026 — 0.027] | -0.604 [-0.609 — -0.598] |
| *R. × bohemica* | Japan | 1 | - | - | - |
|  | Europe | 8 | 0.043 [0.043 — 0.043] | 0.028 [0.028 — 0.028] | -0.551 [-0.553 — -0.548] |
|  | UK | 3 | 0.058 | 0.036 | -0.581 |
|  | North America | 10 | 0.044 [0.043 — 0.044] | 0.028 [0.028 — 0.028] | -0.549 [-0.552 — -0.545] |
| *R. japonica* var. *compacta* | Japan | 5 | 0.053 | 0.040 | -0.341 |
|  | Europe | 2 | 0.058 | 0.037 | -0.560 |
|  | North America | 2 | 0.049 | 0.028 | -0.729 |
| *R. japonica* f. *rosea* | North America | 2 | 0.046 | 0.027 | -0.700 |
| *R. sachalinensis* | Japan | 5 | 0.040 | 0.050 | 0.156 |
|  | Europe | 9 | 0.046 [0.045 — 0.046] | 0.032 [0.032 — 0.032] | -0.506 [-0.510 — -0.501 ] |
|  | North America | 9 | 0.046 [0.046 — 0.046] | 0.033 [0.032 — 0.033] | -0.497 [-0.500 — -0.494] |

### Regional allele pools

When we examined the overlap between regional allele pools of *R. japonica*, we found that the two introduced ranges Europe and North America contained relatively few private alleles but shared a substantial number of alleles with each other (Jaccard similarity index: 0.762) (Fig. 4A, Table S3), whereas the two native regions Japan and China were much more differentiated, with a low similarity index of 0.287. Out of the two native ranges, Japan shared a larger portion of alleles with the introduced ranges (Jaccard similarities: Europe: 0.603, North America: 0.654) than China (Europe: 0.308, North America: 0.325). Out of all regions, Chinese specimens of *R. japonica* had the largest number of private alleles (Fig. 3A). Consistent with the genetic diversity metrics above, the total numbers of alleles were lower in Europe (2,744 alleles) and North America (2,620 alleles) than in Japan (3,063 alleles) and China (3,647 alleles).

**Figure 4.**
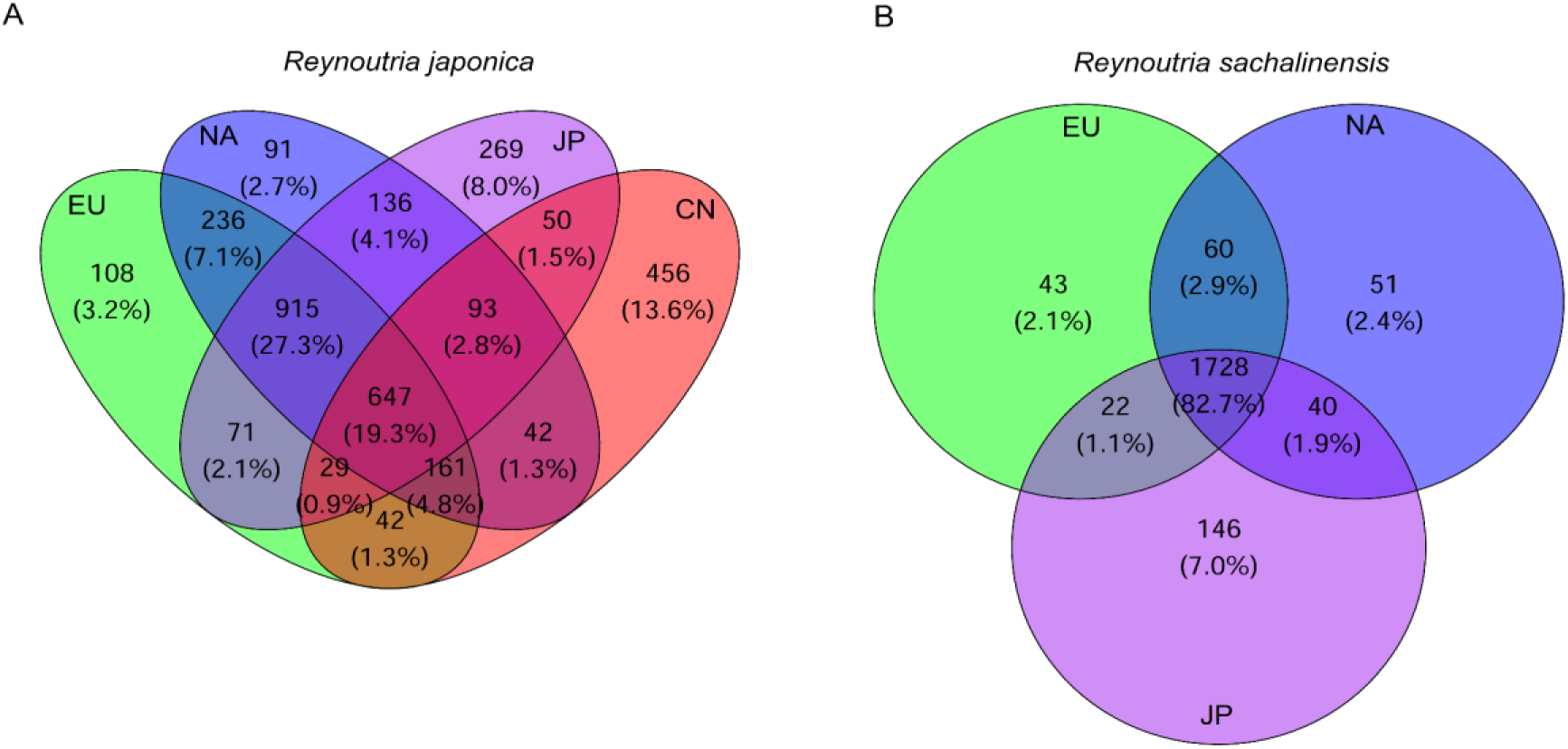
Venn diagrams illustrating the distribution of private alleles (non-overlapping regions) and shared alleles (two-, three-, and four-way intersections) for (A) *R. japonica* across its introduced (EU - Europe, NA- North America) and native range (CN - China, JP- Japan) and (B) *R. sachalinensis* across its native (Japan) and introduced ranges (Europe and North America).

In contrast to *R. japonica*, in *R. sachalinensis* the vast majority of alleles (>80%) were shared across Japan, Europe and North America (Fig. 4B), and the Jaccard similarity indices between all three regions were >0.850 (Table S4). Although the Japanese *R. sachalinensis* samples harbored about three times more private alleles than those from the two introduced regions, the total numbers of alleles were rather similar (∼2,400) in all three regions.

### Phylogenetic relationships

Our phylogenetic analysis based on 1,964 error filtered SNPs identified three well-supported clades (Fig. 5). Clade A primarily comprises the invasive lineage of *Reynoutria japonica* from Europe and North America, a small number of *R. japonica* specimens from Japan representing putative native source populations of the invasion, and the hybrid *R.* × *bohemica* from continental Europe and North America.

**Figure 5.**
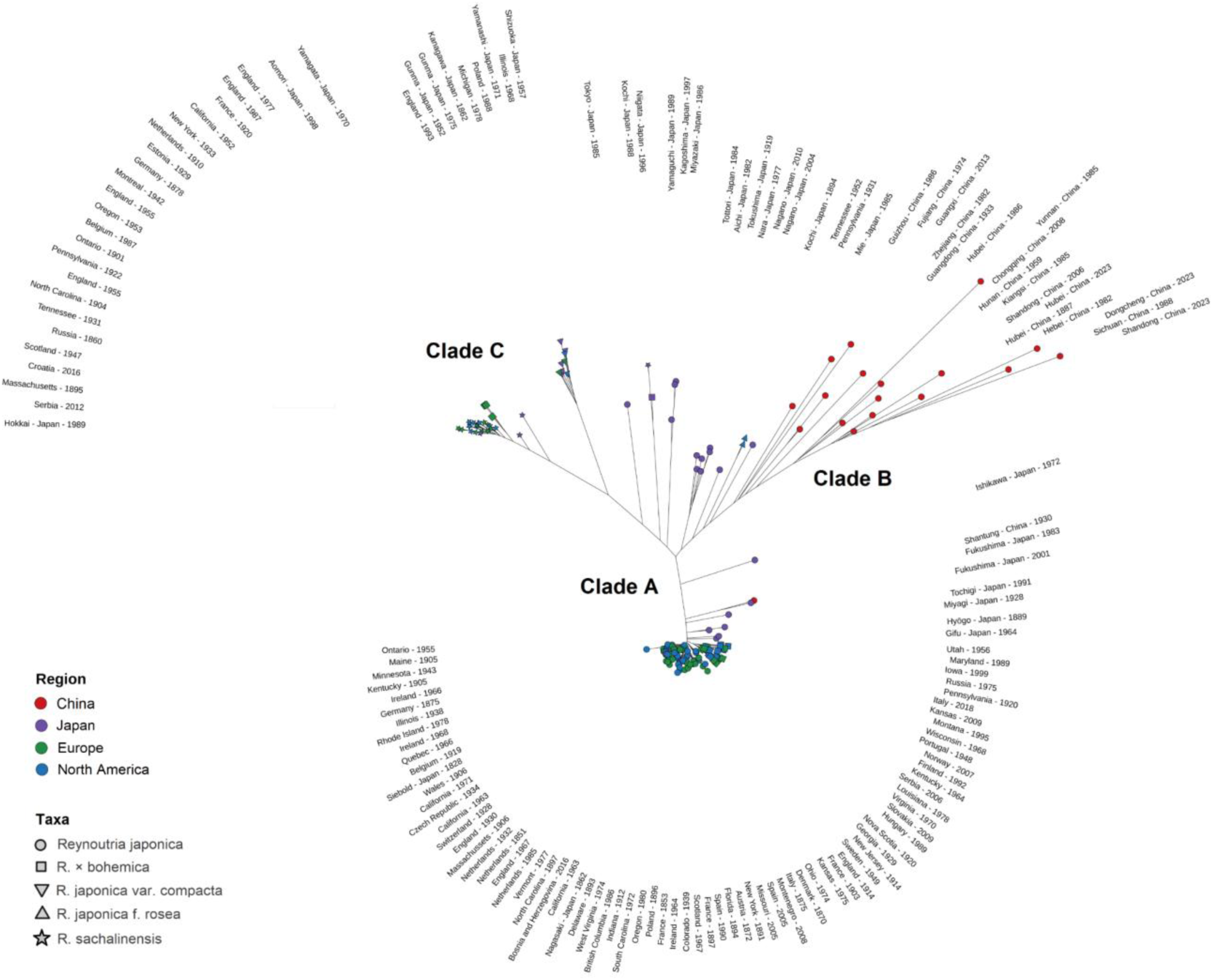
Maximum likelihood phylogenetic tree of Japanese knotweed *s*.*l*. based on 1,964 SNPs from 150 samples. The colours indicate geographic regions, and taxa are indicated by distinct symbols. The tip labels of the phylogenetic tree indicate the collection location and collection year of each specimen.

The clade forms a large cluster with short internal branches, indicating low genetic divergence and recent common ancestry. European and North American samples of *R*. *japonica* are broadly intermixed and do not form distinct regional groups or any other forms of clear geographic structure within and between the two introduced ranges. Likewise, samples collected over a wide temporal span (1828-2016) are distributed throughout the clade without forming basal or derived lineages, indicating no temporal structure. Among the Japanese accessions within Clade A, a specimen collected by Philipp von Siebold in 1828—who is widely credited with introducing Japanese knotweed to Europe—and a second specimen from Nagasaki collected in 1862 both fall within the invasive cluster. Seven additional Japanese samples of *R. japonica* from Tochigi, Miyagi, Hyōgo, Gifu and Fukushima occur on slightly longer branches close to the invasive cluster, indicating that the introduced lineage is part of a regional genetic cluster of Japanese populations. All *R.* × *bohemica* samples from Europe and North America are nested within the invasive *R. japonica* cluster rather than forming a distinct clade. Finally, a single sample from China clusters with a Japanese specimen on a short branch separate from the main invasive cluster, suggesting a recent introduction from Japan to China.

Clade B is composed primarily of *R. japonica* from Asia, including nearly all Chinese samples, eight *R*. *japonica* samples from Japan and two *R. japonica* f. *rosea* individuals from North America. The clade is characterized by long internal branches and spread-out tips. At its base, a cluster of Japanese specimens from central Japan forms shorter branches, indicating closer relatedness, while two additional Japanese samples from southeastern Japan form separate branches. The two North American *R. japonica* f. *rosea* samples branch independently from both the basal Japanese cluster and the Chinese accessions, consistent with independent introductions or a distinct cultivated variety. Chinese specimens are distributed across long, divergent branches, reflecting historical separation from Japanese populations. Similar to clade A, no clear temporal structure is observed (Fig. 5).

Clade C is the most heterogeneous clade in the phylogeny, comprising multiple taxa and geographic regions. It includes *R. sachalinensis* from northern Japan, the Russian Far East (*e.g*., Sakhalin), Europe and North America, as well as three *R.* × *bohemica* samples from the UK and France. A secondary branch contains *R. japonica* var. *compacta* from Japan and introduced regions (Europe and North America). In addition, three distinct branches are observed: one containing a Japanese *R. japonica* var. *hachidyoensis* sample; another including a Japanese *R.* × *bohemica* sample together with a *R. sachalinensis* specimen from southwestern Japan and a third comprising three *R. japonica* specimens from southern Japan. This clade is characterized by long internal and basal branches but also numerous short terminal branches particularly among *R. sachalinensis* samples (Fig. 5).

## Discussion

Genetic studies of introduced populations help to reveal the evolutionary processes underlying initial species establishment and subsequent colonization. Although invasive species often experience reduced genetic diversity due to bottlenecks and founder effects, multiple introduction events can counteract these losses by increasing genetic variation and promoting recombination among source genotypes—potentially contributing to invasion success. However, reconstructing introduction histories is challenging because the full range of genotypes that had the opportunity to colonize is rarely known. In this study, we address important gaps in the invasion history of Japanese knotweed *s*.*l*. by incorporating extensive temporal sampling in the native range in Japan and China and including the earliest documented herbarium specimens from the introduced ranges in Europe (Leiden: 1851) and North America (New York: 1891).

### Multi-locus lineages

We identified a relatively small number of multi-locus lineages (MLLs) within this species complex, consistent with the predominantly clonal nature of these taxa. Japan comprised six MLLs along with an additional unique genotype corresponding to *R. japonica* var. *hachidyoensis*. Two MLLs were restricted to Japan: one comprising *R. japonica* from the southernmost region (Kyushu), and another grouping a *R. sachalinensis* specimen from southwestern Japan with a *R.* × *bohemica* individual from western Japan. The remaining Japanese MLLs were also detected in other regions. These included (i) the invasive *R. japonica* lineage present in western Europe and North America; (ii) a second *R. japonica* lineage from central and southern Japan shared with China and North America; and (iii) lineages corresponding to *R. japonica* var. *compacta* (one lineage) and *R. sachalinensis* (one lineage). North America contained four MLLs: three shared with Europe and Japan, and one corresponding to *R. japonica* f. *rosea* which was shared only with Japan and China. Europe harbored three MLLs, all of which were also present in North America and Japan. China comprised one widespread *R. japonica* lineage shared with Japan and North America, and a second lineage restricted to northeastern China that belonged to the invasive *R. japonica* lineage found across Europe, North America and Japan. It is important to note that using a single genetic distance threshold across different taxa, combined with a reference genome that corresponds to only one species in the genus, may have underestimated MLL diversity—particularly in *R. sachalinensis* and the hybrid *R.* × *bohemica* (Stafiniak et al., 2025). Current efforts are underway to sequence and assemble the genome of *R. sachalinensis* by the Darwin Tree of Life Project Consortium (Data Portal, accessed 02 February 2026), which promises to provide additional insights into the genomics and taxonomy of this polyploid species complex.

Previous studies assessing clonality in this species complex have yielded conflicting results. Using genotyping-by-sequencing (GBS) and a Euclidean-distance threshold to delineate genotypes, VanWallendael et al. (2020) identified as many as 66 multilocus genotypes (MLGs) in *R. japonica*, along with high levels of clonality in *R. sachalinensis* and *R.* × *bohemica* across eastern North America. Similarly, Grimsby et al. (2007), employed seven microsatellite markers and a Simpson’s diversity index incorporating ramet density, and detected 26 MLGs among *R. japonica sensu lato* samples from three populations in Massachusetts. Conversely, Gaskin et al. (2014) identified a single *R. japonica* genotype across western North America but multiple distinct genotypes of *R.* × *bohemica* and *R. sachalinensis* using AFLP markers. These discrepancies likely reflect both limited marker resolution and the specific error threshold used to genotype the individuals. To mitigate the influence of sequencing errors on MLL assignment, we included 11 samples, each analyzed in three technical replicates. However, our inferences were limited by the characteristics of the dataset, particularly low sequencing coverage (mean coverage of 19× across samples). This precluded estimation of somatic mutation accumulation (Klimešová et al., 2025). While assessing ploidy would have provided further insight, low coverage sequencing and degraded herbarium DNA precluded flow cytometry. Overall, the observed pattern of limited MLL diversity is consistent with patterns reported in other clonally reproducing organisms. For example, cassava cultivation has led to an almost complete shift toward clonal propagation globally (Kistler et al., 2025), while several seagrass species, including *Zostera marina* and *Posidonia oceanica*, exhibit extensive population-level clonality (Yu et al., 2020; Arnaud-Haond et al., 2012). Similarly, global monoclonality has been documented in *Pennisetum setaceum* following the spread of a single “super-genotype” (Le Roux et al., 2007).

### Genetic diversity

Both *R. japonica* and *R*. *sachalinensis* in Europe and the US showed marked reductions in nucleotide diversity relative to their native range, whereas observed heterozygosity only showed moderate or no decline. FIS values shifted from near zero in China, indicating random mating, to moderately positive in Japan, suggesting some inbreeding, and were strongly negative in the introduced range, consistent with predominantly clonal reproduction. *Reynoutria japonica* var. *compacta* exhibited a pattern similar to *R. japonica*, but with negative FIS values in both native and introduced ranges. In the introduced range, *R*eynoutria × *bohemica* exhibited comparable or lower reductions in genetic diversity relative to both *R. japonica* and *R. sachalinensis*. These patterns of genetic diversity align with broader observations in other introduced species. A meta-analysis spanning 80 species (including animals, plants and fungi) demonstrated significant reductions in allelic richness and heterozygosity in introduced populations relative to their sources (Dlugosch & Parker 2008), consistent with diversity loss during founding events (but see Rollins et al., 2013; Bhattacharya et al., 2022 for exceptions). For *R. japonica*, both total and private alleles were also higher in the native range than in the introduced range, while *R. sachalinensis* showed similar levels across ranges. In the native range, we identified a considerable number of private alleles (*R. japonica*: China: 456, Japan: 269; *R. sachalinensis*: Japan: 146), suggesting that the introduced populations did not capture the full genetic diversity present in the native sources. Interestingly, we also observed some private alleles in both *R. japonica* and *R. sachalinensis* in Europe and North America. This may indicate either gaps in our sampling of native regions that could have contributed to these alleles, new mutations, the influence of paralogs and orthologs or simply background noise. Notably, the majority of the private alleles in the introduced range were singletons, whereas native populations had much lower proportions of singletons. This indicates that introduced populations harbor very little new genetic variation and that most observed differences likely reflect rare variants or sequencing artifacts, thereby rendering temporal regressions analyses largely uninformative.

These diversity patterns must be interpreted in light of the species’ genomic architecture. *Reynoutria japonica* is a mixed auto-allooctoploid (AAAABBBB) originating from a genome duplication of *R. sachalinensis*. One subgenome (A) shares a common ancestor with the diploid *Fallopia multiflora* (Wang et al., 2025a). This characteristic may enable *R*. *japonica* to retain high levels of non-segregating “fixed heterozygosity” from divergent parental genomes (Soltis & Soltis, 2000). *Reynoutria sachalinensis* is allotetraploid (AABB), and *R.* × *bohemica* is an allohexaploid (AAABBB) resulting from hybridization between *R. japonica* and *R. sachalinensis* (Wang et al., 2025a). Although *R. japonica*, *R. sachalinensis*, and *R.* × *bohemica* share the same A and B subgenomes, differences in ploidy and sequence divergence may underestimate variation in divergent or hybrid regions when mapping to the *R. japonica* reference genome (Van Wallendael et al., 2021; Stafiniak et al., 2025). Reference-free approaches, such as k-mer-based analyses, are likewise constrained by the low sequencing coverage of this dataset. Consequently, our results should be viewed as conservative estimates of genetic diversity rather than measures of absolute species-wide diversity.

### Evolutionary relationships, patterns of introduction and hybridization

We found strong support that the introduction of *R. japonica* into Europe and North America traces back to a single multi-locus lineage that occurs widely across Japan, with matches to native accessions from Nagasaki (where botanical collections were made by Philipp von Siebold), as well as Hyōgo (west), Gifu (center) and northeast: Miyagi, Fukushima and Tochigi. This is consistent with previous chloroplast-based studies, which identified a dominant Japanese haplotype shared across Europe and North America (Pashley 2003; Zhang et al., 2024, and confirmed by our study, see Fig. S1). Importantly, this genetic evidence for a Japanese origin is mirrored by phenotypic patterns observed in experimental studies. Common garden experiments comparing introduced *R. japonica* populations from the US, Europe and the native range (China and Japan) demonstrated that introduced populations differ from those in China but exhibit similar trait values to plants from Japan including higher clonality and greater plasticity in most traits (Cao et al., 2025; Wang et al., 2025b). The near-identical genetic composition of North American and European *R. japonica* individuals combined with extensive allele sharing suggests that western Europe acted as a bridgehead, facilitating the subsequent introduction and spread of this species in North America. The transfer of this same *R. japonica* lineage from Europe to North America was likely facilitated by horticultural exchange, as documented by del Tredici (2017) and is consistent with previous genetic studies (Gammon & Kesseli, 2010; Gaskin et al., 2014; Richards et al., 2012; VanWallendael et al., 2021, Zhang et al., 2024) and archival records (Bailey & Conolly 2000).

Interestingly, in contrast to previous studies that reported no shared *R. japonica* lineages between the introduced ranges and China (Pashley 2003; Zhang et al., 2024), we found a Chinese accession (36°N, Tsingtao, collected in 1930) that clustered within the invasive lineage, diverging on the same branch as a Fukushima (Japan) accession. This suggests that some northeastern Chinese populations may have been recently introduced from Japan (Zhang et al., 2024). *Reynoutria japonica* has long been used as a food and medicinal plant in China and Japan (Shimoda & Yamasaki, 2016) and is also planted for roadside erosion control (Pashley 2003), practices that likely promoted its spread well beyond what natural dispersal would allow.

In addition to the widespread invasive lineage of *R. japonica*, we detected independent introductions of *R. japonica* var*. compacta* from Japan into Europe and North America, as well as introductions of the horticultural garden cultivar *R. japonica* f. *rosea* into North America. However, neither taxon appears to have substantially contributed to the invasion to date, as they are only rarely observed (Bailey & Conolly 2000; Forman & Kesseli 2003; Jovanović et al., 2018), but they could still serve as a pollen source for the widespread *R. japonica* lineage. As a result, previous reports of multiple introduction events of distinct *R. japonica* genotypes in North America (del Tredici 2017; Gammon & Kesseli 2010) may, in fact, pertain to *R. japonica* var. *compacta* and the garden cultivar.

Regarding the other parental taxa, all introduced *R. sachalinensis* samples formed a single group in both the combined and chloroplast phylogenies, showing genetic similarities with native populations from northern Japan and southern Sakhalin Island, a pattern that aligns with historical records and horticultural spread via nurseries and botanical gardens (Bailey & Conolly 2000). The high genetic and allelic overlap between the introduced regions (Europe and North America) indicates that the introduction process in these areas involved common founder populations, leading to similar genetic profiles.

We identified two distinct *R.* × *bohemica* hybrid lineages in the introduced range. The widespread lineage in Europe and North America exhibited a multilocus genetic profile closely matching *R*. *japonica*, suggesting backcrossing or later-generation hybrids—a pattern that may contribute to their invasive success if these hybrids predominantly express *R. japonica* traits. These hybrids are genetically diverged from introduced *R*. *sachalinensis*, the putative male parent, which may reflect either a “dilution” of the *R*. *sachalinensis* genome through backcrossing or an artifact of using the *R*. *japonica* genome as a reference for SNP calling (VanWallendael et al., 2021). Such hybridization can strongly influence the genomic composition of *R*. × *bohemica*, with potential ecological consequences. Previous studies have shown that although *R. japonica* invades a broader range of habitat types than both *R. sachalinensis* and *R.* × *bohemica*, the hybrid outcompetes the parental species at sites where the taxa co-occur (Bímová et al., 2004). Moreover, *R.* × *bohemica* exhibits novel niche adaptations that are not merely intermediate between those of its parental species. These adaptations facilitate its range expansion in southeastern Europe and the Mediterranean while also conferring enhanced drought tolerance (Bailey & Wisskirchen 2004; Jovanović et al., 2018). The second multi-locus hybrid lineage, comprising three hybrids (two from England, one from France), was positioned in the phylogeny between *R*. *sachalinensis* and *R*. *japonica* var. *compacta*. These hybrids carried the *R*. *sachalinensis* plastid and showed reduced NEI genetic divergence from *R*. *sachalinensis*. Because both parental taxa occur as male and female plants in the wild, hybridization can occur in either direction. One British hybrid was reported on the label with 44 chromosomes, suggesting that the other two are likely also tetraploids, consistent with the higher prevalence of tetraploid *R*. × *bohemica* and *R*. *japonica* var. *compacta* in the British Isles compared to continental Europe (Bailey & Wisskirchen 2004). Given that both parental taxa show limited invasive potential, this hybrid lineage may pose a lower ecological risk and remain geographically restricted. Nevertheless, our data does not allow us to determine whether these hybrids originated *in situ* or were introduced via horticultural seed distribution (see also Hollingsworth & Bailey 2000b). In our sampling, only a single Japanese accession (collected in 1996 from the northwest Honshu) displayed the characteristic *R.* × *bohemica* morphology. In the phylogeny, this sample diverged alongside a *R. sachalinensis* accession from Shikoku while both clustered with southern Japanese *R. japonica* in the plastid tree, suggesting *R. sachalinensis* cpDNA capture via introgression (Pashley 2003; Park et al., 2018). In the native range, the hybrid—recently described as *Reynoutria* × *mizushimae* Yokouchi ex T. Shimizu (Tatemi 1997)—occurs primarily where *R. sachalinensis* and *R. japonica* co-occur in lowland northern Honshu (Aomori prefecture) but is rarer elsewhere (Pashley 2003). Hybridization between the two taxa has been reported sporadically in Korea (Kim & Park 2000), and field surveys in New Zealand indicate both direct introduction of *R.* × *bohemica* from Japan as well as *in situ* hybrid formation (Desjardins et al., 2023). Further sampling and genetic analyses across native and introduced ranges are needed to better understand the diversity and origins of hybrid lineages.

Our study also provides insights into the genetic structure of native *R. japonica* populations. Chinese populations were genetically distinct from all other regions, exhibiting the highest levels of genetic diversity and a wide range of plastid types, yet they contained only two multi-locus lineages and did not contribute to the invasion. The presence of different maternal lineages based on plastid types could reflect historical events (such as glaciation), with multiple plastid types evolving in isolation, but without the diversification of multi-locus genetic lineages. Overall, Chinese populations appear to have diverged from *R*. *japonica* populations in southern Japan, which aligns with a previous study that used chloroplast markers and ancestral state reconstruction (Zhang et al., 2024). Alternatively, this pattern could also result from limitations in our methodology, which may have missed finer intraspecific genetic variation. Japan also displayed some spatially structured genetic variation. A field study in Japan (Pashley 2003) found that the predominant cytotype of *R*. *japonica* was tetraploid, with octoploids being less common. These cytotypes appear to exhibit distinct ecological preferences: tetraploids are more abundant on the east coast, characterized by hot, humid summers and dry winters, while octoploids are more frequent on the west coast, which experiences heavy winter snow and cooler summers (Ohsawa et al., 2017). Although originally introduced for ornamental purposes (del Tredici, 2017), the octoploid lineage that established in Europe and North America (Irimia et al., 2025) may have inherent ecological and physiological traits that facilitated its invasive success.

High clonality seems to be prevalent across the introduced range of Japanese knotweed *s.l*. Our results confirm that a single multi-locus lineage of *R. japonica*, originating in Japan, was introduced to Europe and North America. This lineage’s success in colonizing a wide range of habitats may be due to a combination of traits, high phenotypic plasticity and ecological advantages conferred by polyploidy (Mounger et al., 2021; Cao et al., 2025; Irimia et al., 2025; Wang et al., 2025b). Our study also highlights the complexity of hybridization events in the invasion process. To better guide management efforts, future research should focus on exploring the diversity of hybrid lineages and their roles in the ongoing spread of this invasive species.

## Acknowledgments

We thank the curators of the following herbaria: ASU, BRNU, C, CAN, CAS, CFSI, CGE, CM, DOV, E, FI, FLAS, G, GA, GB, GJO, GMNJ, HYO, IBK, ILL, ILLS, IMC, KANU, KRAM, LD, LY, M, MA, MBK, MIN, MOR, MT, N, NCU, NCY, NMU, NY, OXF, P, PE, RSA, TAA, TENN, TROM, TUR, UPS, US, UT, WU and ZA for providing us plant material for genetic analyses. We are grateful to Carolina Royo Brun (ICVV, Spain) for her assistance with the DNA isolation protocol.

## Data availability

The raw sequencing data generated in this study will be deposited in the NCBI Sequence Read Archive (SRA) and will be made publicly available upon acceptance of the manuscript for publication.

