## Supplementary Material for "Using herbarium genomics to understand the history of a global plant invasion"

**Table S1**. Overview of the herbarium samples used in this study, with sample identifiers (barcodes or catalog numbers) collector names and collection numbers, the original taxon identification as recorded by the herbarium collectors, the collection year and institution or source herbarium (herbarium code based on Index Herbariorum, see: <https://sweetgum.nybg.org/science/ih/>), the phenological stage of specimens (vegetative, unopened flower buds, female or male flowers or fruiting), further information about the sampling origin (area of origin, habitat, geographic coordinates). The last three columns provide the assignments to multi-locus lineages (MLLs) and taxa identity in our study (RJA = *R*. *japonica*, RBO = *R*. × *bohemica*, RSA = *R*. *sachalinensis*, RJC = *R*. *japonica* var. *compacta*, RJR = *R*. *japonica* *f*. *rosea*, RJU = *R*. *japonica var. uzenensis* Honda, RJH = *R*. *japonica* var. *hachidyoensis* (Makino) Nakai; RJFC = *R*. *japonica* *f*. *colorans* Makino) and the DNA library preparation method (ds_half-UDG = double stranded library, partially treated with uracil DNA glycosylase; ds_non-UDG = without uracil DNA glycosylase treatment). Samples with asterisks after the barcode were technical replicates.

| **Catalog/Barcode** | **Collector & CollectionNo** | **Taxon** | **Year** | **Institution** | **Phenology** | **State/Country/Province** | **Habitat** | **Lat** | **Lon** | **MLL** | **Taxon**  **assignment** | **Library**  **prep method** |
| --- | --- | --- | --- | --- | --- | --- | --- | --- | --- | --- | --- | --- |
| BEOU46519* | D. Lakušić, N. Kuzmanović & I. Janković 46519 | RJA | 2016 | BEOU | ♀ flowers | Bosnia and Herzegovina | village, ruderal | 44.3777 | 17.9932 | 3 | RJA | ds_half-UDG |
| 60886 | V. Šegota | RSA | 2016 | ZA | vegetative | Croatia | riverbank | 45.5942 | 14.6491 | 4 | RSA | ds_half-UDG |
| FI058969 | N. Hofmann | RBO | 2018 | FI | ♂ flowers | Italy | city, road embankment | 43.7305 | 12.6358 | 3 | RBO | ds_half-UDG |
| BEOU27311 | D. Lakušić, B. Lakušić & M. Praščević 27311 | RJA | 2008 | BEOU | vegetative | Montenegro | village, cultivated | 42.6184 | 19.9238 | 3 | RJA | ds_half-UDG |
| V-174556 | B.G. Østerkløft 122 | RJA | 2007 | TROM | vegetative | Norway | city, scrap soil area | 67.3206 | 14.4604 | 3 | RBO | ds_half-UDG |
| BEOU37718 | S. Jovanović, D. Lakušić, J. Šinžar-Sekulić 37718 | RSA | 2012 | BEOU | fruits | Serbia | village, pond embankment | 45.3060 | 20.4298 | 4 | RSA | ds_half-UDG |
| GJO_0088103 | V. Mikoláš 2619/09 | RBO | 2009 | GJO | fruits | Slovakia | village | 48.7188 | 21.1826 | 3 | RBO | ds_half-UDG |
| MA782918 | S. Castroviejo & V.R. Gracia 17814 | RJA | 2005 | MA | ♀ flowers | Spain | highway ditch | 42.73 | -7.32703 | 3 | RJA | ds_half-UDG |
| BR000001229335 | J. Saintenoy-Simon | RSA | 1987 | BR | vegetative | Belgium | roadside | 50.3602 | 4.86807 | 4 | RSA | ds_half-UDG |
| P05000007 | D.E. Boufford & B. Bartholomew 24101 | RJA | 1988 | P | ♀ flowers | Sichuan / China | mountain range, at gorge, remnant forest | 30.99468 | 103.5227 | 1 | RJA | ds_half-UDG |
| P04961759 | E. Cosson | RJA | 1897 | P | ♀ flowers | France | city, private garden | 45.6403 | 0.13974 | 3 | RJA | ds_half-UDG |
| B 10 1201971 | F.A. Forhering | RJA | 1875 | B | fruits | Germany | city, in garden | 49.3987 | 8.67243 | 3 | RJA | ds_half-UDG |
| B 10 0481650 | Herbarium Erzberger | RSA | 1989 | B | unopened flowers | Hungary | city | 47.5096 | 18.9712 | 3 | RBO | ds_half-UDG |
| L. 0768025 | P. Florusse & J.W. Jongepier 708 | RJA | 1985 | L | ♂ flowers | Netherlands | city, railway yard | 51.9420 | 4.48144 | 3 | RJA | ds_half-UDG |
| M-0323153 | V.V. Makarov | RJA | 1981 | M | ♀ flowers | Russia | vacant lot | 55.8440 | 37.5864 | 3 | RJA | ds_half-UDG |
| Z-000234653 | E. Müller | RJA | 1928 | Z | ♀ flowers | Switzerland | village | 47.1240 | 9.31220 | 3 | RJA | ds_half-UDG |
| CGE00070939 | M. MCallon Webster 11,620 | RJA | 1967 | CGE | ♀ flowers | Scotland / UK | abandoned sawmill | 57.5118 | -3.97789 | 3 | RJA | ds_half-UDG |
| LY0005293 | R. Bonaparte | RJA | 1903 | LY | ♀ flowers | France | city, close to restaurant | 48.8678 | 2.31231 | 3 | RJA | ds_half-UDG |
| B 10 1201981 | R. Oldham 699 | RJA | 1862 | B | unopened flowers | Kyushu / Japan | city | 32.7502 | 129.877 | 3 | RJA | ds_half-UDG |
| B 10 1201970 | R. Gross | RJA | 1896 | B | ♀ flowers | Poland | city, ruderal | 54.2080 | 19.1175 | 3 | RJA | ds_half-UDG |
| G00421221* | P. Silva, Fontes & Rainha 2539 | RJA | 1948 | G | ♂ flowers | Portugal | mountain range, field edge | 41.8250 | -8.04105 | 3 | RBO | ds_half-UDG |
| TFA.567401 | E. Piispala 15/92 | RBO | 1992 | TUR | vegetative | Finland | city, edge of parking area | 63.0855 | 21.6201 | 3 | RBO | ds_half-UDG |
| 03759204 | C.Y. Chiao 2506 | RJA | 1930 | NY | ♀ flowers | Shandong / China | city park, sandy slope | 36.0605 | 120.356 | 3 | RJA | ds_half-UDG |
| ILL00129901 | O.W. Knight | RJA | 1905 | ILL | unopened flowers | Maine / USA | city, roadside | 44.8929 | -68.6933 | 3 | RJA | ds_half-UDG |
| RSA0503144 | W.H. Aiken | RJA | 1905 | RSA | ♀ flowers | Kentucky / USA | city | 39.0858 | -84.4923 | 3 | RJA | ds_half-UDG |
| RSA0503141* | T.C. Fuller 11152 | RSA | 1963 | RSA | ♀flowers | California / USA | roadside, on highway | 41.2713 | -122.272 | 3 | RJA | ds_half-UDG |
| RSA0503137 | H.E. & S.T. Parks 24448 | RSA | 1952 | RSA | ♂flowers | California / USA | roadside, on highway | 40.7553 | -124.191 | 4 | RSA | ds_half-UDG |
| FLAS 8116 | P.N. Rolfs | RJA | 1894 | FLAS | unopened flowers | Florida / USA | city, cultivated | 30.1874 | -82.6411 | 3 | RJA | ds_half-UDG |
| FLAS 31625 | G.F. Weber | RJA | 1939 | FLAS | ♀flowers | Colorado / USA | city, cultivated | 40.5628 | -105.068 | 3 | RJA | ds_half-UDG |
| 037781 | O.E. Jennings & L.C. Denise | RJA | 1920 | CM | fruits | Pennsylvania / USA | city | 40.5097 | -80.0981 | 3 | RBO | ds_half-UDG |
| ILLS00078641 | J.C. Carter 8150 | RJA | 1938 | ILLS | vegetative | Illinois / USA | near village | 41.8003 | -87.9365 | 3 | RJA | ds_half-UDG |
| 03262258 | - | RSA | 1895 | US | ♀ flowers | Massachusetts / USA | city, cultivated | 42.3726 | -71.1122 | 4 | RSA | ds_half-UDG |
| 0084518MOR* | G. Murata 45633 | RJA | 1985 | MOR | unopened flowers | Kyushu / Japan | mountain foot, on grassy slope | 33.2820 | 131.390 | 2 | RJA | ds_half-UDG |
| KANU351723 | C.C. Freeman 21251 & C. Morse | RJA | 2005 | KANU | ♀ flowers | Missouri / USA | city, wooded lot along the street | 39.1881 | -93.8834 | 3 | RJA | ds_half-UDG |
| GA140493 | J.H. Pyron E6722 | RJA | 1929 | GA | ♀flowers | Georgia / USA | cultivated, old botanical garden | 33.9021 | -83.3848 | 3 | RJA | ds_half-UDG |
| KANU00082404 | R. Brooks 11861 | RJA | 1975 | KANU | fruits | Kansas / USA | in ditch, at roadside | 39.1759 | -94.7710 | 3 | RJA | ds_half-UDG |
| 1085679 | O. Laheln 5508 | RJA | 1943 | MIN | unopened flowers | Minnesota / USA | creek woods | 46.8262 | -92.0741 | 3 | RJA | ds_half-UDG |
| TENN-V-0087579 | R.L. James 17110 | RJA | 1952 | TENN | fruits | Tennessee / USA | city, cultivated | 36.3357 | -82.3471 | 1 | RJR | ds_half-UDG |
| TENN-V-0088577 | C.L. Rodgers & N.E. Mullens 0253 | RJA | 1972 | TENN | unopened flowers | South Carolina / USA | country road, rest area | 34.8708 | -82.3797 | 3 | RJA | ds_half-UDG |
| TENN-V-0088557 | R.D. Thomas 60668 | RJA | 1978 | TENN | unopened flowers | Louisiana / USA | near creek, south of village | 32.1519 | -92.9800 | 3 | RBO | ds_half-UDG |
| ILL00129908 | R.R. Halse 2282 | RJA | 1980 | ILL | ♀ flowers | Oregon / USA | gravel pits | 45.0295 | -123.054 | 3 | RJA | ds_half-UDG |
| 0084512MOR | E.F. Evert 29724 | RJA | 1995 | MOR | vegetative | Montana / USA | disturbed riparian area | 45.7303 | -109.662 | 3 | RBO | ds_half-UDG |
| UT0073333 | W.P. Cottam | RJA | 1956 | UT | unopened flowers | Utah / USA | escape, in a field | 40.7857 | -111.889 | 3 | RBO | ds_half-UDG |
| UT0073288 | S.H. Sohmer 5107f | RJA | 1968 | UT | ♂ flowers | Wisconsin / USA | city, behind cemetery | 43.8206 | -91.2333 | 3 | RBO | ds_half-UDG |
| TOYA107657 | Y. Nagai | RJA | 1972 | TOYA | vegetative | Honshu / Japan | sea coast | 37.1500 | 136.983 | 3 | RJA | ds_half-UDG |
| TOYA86770 | M. Ohta | RJA | 1985 | TOYA | ♂ flowers | Honshu / Japan | city | 33.9633 | 136.107 | 1 | RJA | ds_half-UDG |
| TOYA86153 | M. Ohta | RSA | 1989 | TOYA | ♂ flowers | Hokkaido / Japan | village, at bridge | 42.9941 | 142.347 | 4 | RSA | ds_half-UDG |
| TOYA84947 | M. Ohta | RJA | 1986 | TOYA | ♀ flowers | Kyushu / Japan | city | 31.7575 | 131.464 | 2 | RJA | ds_half-UDG |
| TOYA83438 | M. Ohta | RBO | 1996 | TOYA | vegetative | Honshu / Japan | forest road | 37.9499 | 138.500 | 5 | RBO | ds_half-UDG |
| TOYA55320 | M. Ohta | RJA | 1984 | TOYA | fruits | Honshu / Japan | city | 35.3862 | 133.485 | 1 | RJA | ds_half-UDG |
| TOYA42355 | K. Uenohara 306 | RJA | 1982 | TOYA | vegetative | Honshu / Japan | town | 34.8641 | 137.297 | 1 | RJA | ds_half-UDG |
| C1-080297 | M. Hashimoto 1534 | RJU | 1964 | HYO | fruits | Honshu / Japan | city | 35.4656 | 136.619 | 3 | RJA | ds_half-UDG |
| B 10 1201980 | C.J. Maximowicz | RJA | 1862 | B | unopened flowers | Honshu / Japan | city | 35.4437 | 139.638 | 6 | RJA | ds_half-UDG |
| P05000037* | B.P. Bodinier 294 | RJA | 1889 | P | fruits | Honshu / Japan | hills close to waterfall | 34.7079 | 135.179 | 3 | RJA | ds_half-UDG |
| MBK0292730 | A. Seo 1447 | RJA | 1997 | MBK | fruits | Kyushu / Japan | city | 31.2125 | 130.646 | 2 | RJC | ds_half-UDG |
| FKSE 69323 | Y. Komine, M. Yoshikawa, M. Yaita | RJA | 1991 | FKSE | fruits | Honshu / Japan | in the valley, near the bridge | 36.7964 | 139.817 | 3 | RJA | ds_half-UDG |
| FKSE 7311 | H. Sase | RJH | 1985 | FKSE | fruits | Honshu / Japan | city | 34.7378 | 139.394 | NA | RJH? | ds_half-UDG |
| B 10 1201978 | B. Krug 680 | RJA | 1919 | B | fruits | Shikoku / Japan | city | 34.0686 | 134.542 | 1 | RJA | ds_half-UDG |
| CM297915 | T. Nemoto & H. Hoshi 1682 | RJA | 1983 | CM | fruits | Honshu / Japan | lakeside | 37.4539 | 139.982 | 3 | RJA | ds_half-UDG |
| B 10 1201982 | K. Okamoto 693 | RJA | 1952 | B | fruits | Honshu / Japan | mountain area | 36.2076 | 138.635 | 6 | RJA | ds_half-UDG |
| Z-000234658 | H. Koyama & M. Hotta 5426 | RJA | 1977 | Z | fruits | Honshu / Japan | grassy slope, in light forest | 34.2419 | 135.947 | 1 | RJA | ds_half-UDG |
| WU0131126 | - | RJA | 1894 | WU | ♀ flowers | Shikoku / Japan | city | 33.5665 | 133.130 | 1 | RJR | ds_half-UDG |
| - | E. Ishiba | RSA | 1928 | GB | ♂ flowers | Honshu / Japan | city | 38.2564 | 140.842 | 3 | RJA | ds_half-UDG |
| B 10 1201967 | M. Togasi 1548 | RJC | 1957 | B | ♂ flowers | Honshu / Japan | mountain area | 35.3606 | 138.727 | 6 | RJC | ds_half-UDG |
| B 10 1201963 | H. Ohashi, Y. Tateishi & H. Ohba 708133 | RSA | 1970 | B | unopened flowers | Honshu / Japan | on sunny and moist banks | 38.3093 | 140.475 | 4 | RSA | ds_half-UDG |
| B 10 0426659 | F. Konta & J. Kitagawa 35759 | RJA | 2010 | B | ♂ flowers | Honshu / Japan | at moist place, along conifer plantation | 35.2702 | 137.740 | 1 | RJA | ds_half-UDG |
| CM435363 | J.A. Shafer | RSA | 1901 | CM | fruits | Ontario / Canada | waterfalls | 43.0796 | -79.0871 | 4 | RSA | ds_non-UDG |
| 0084521MOR | G.B. Straley 3794 | RJA | 1986 | MOR | unopened flowers | British Columbia / Canada | entrance to nature trail, university lands | 49.2438 | -123.214 | 3 | RJA | ds_non-UDG |
| B 10 1201979 | A. Henry 2290 | RJA | 1887 | B | unopened flowers | Hubei / China | city | 30.6918 | 111.286 | 1 | RJA | ds_non-UDG |
| TAA0091032 | J. Talts | RSA | 1929 | TAA | vegetative | Estonia | city, seashore | 58.3767 | 24.4881 | 4 | RSA | ds_non-UDG |
| P04555472 | E. Kugler | RSA | 1878 | P | ♀ flowers | Germany | in garden, at the castle | 47.8660 | 9.99458 | 4 | RSA | ds_non-UDG |
| M0323149 | Leuze & Doppelbaur | RJA | 1964 | M | unopened flowers | Ireland | wasteland | 52.1418 | -10.2784 | 3 | RJA | ds_non-UDG |
| FI065948 | B. Hausmann | RJA | 1875 | FI | ♀ flowers | Italy | city, vineyards | 46.4970 | 11.3444 | 3 | RJA | ds_non-UDG |
| LY0005301 | P.F. von Siebold | RJA | 1828 | LY | ♀ flowers | Japan | NA | 32.7631 | 129.839 | 3 | RJA | ds_non-UDG |
| P04940026* | W.H. de Vriese | RJA | 1851 | P | unopened flowers | Netherlands | botanical garden | 52.1571 | 4.48459 | 3 | RJA | ds_non-UDG |
| BEOU55742* | M. Cvetković | RJA | 2006 | BEOU | fruits | Serbia | city, school playground | 44.7718 | 20.4899 | 3 | RBO | ds_non-UDG |
| MA509322 | E. Monasterio & J. Pedrol 4363 | RJA | 1990 | MA | unopened flowers | Spain | roadside | 42.8005 | 0.70369 | 3 | RJA | ds_non-UDG |
| 0128371 | E. Kjellgren | RJA | 1949 | GB | ♀ flowers | Sweden | airfield | 59.4441 | 13.3482 | 3 | RJA | ds_non-UDG |
| NY03759272 | J. Lindley | RJA | 1850 | NY | ♀ flowers | England / UK | in city, private garden | 51.5054 | -0.27683 | 3 | RJA | ds_non-UDG |
| KANU00324585 | C.C. Freeman 12703 | RJA | 1999 | KANU | vegetative | Iowa / USA | ditch, park entrance | 42.8795 | -92.3187 | 3 | RBO | ds_non-UDG |
| KANU363875 | C.A. Morse 20006 | RBO | 2009 | KANU | ♂ flowers | Kansas / USA | roadside, edge of mining pit | 37.4924 | -94.6865 | 3 | RBO | ds_non-UDG |
| 1085708 | J.H. Wibbe | RJA | 1891 | MIN | ♀ flowers | New York / USA | riverbanks | 42.8224 | -73.9392 | 3 | RJA | ds_non-UDG |
| DOV0030381 | W.M. Canby | RJA | 1893 | DOV | ♀ flowers | Delaware / USA | city, waste grounds | 39.7463 | -75.5431 | 3 | RJA | ds_non-UDG |
| ILL00129906* | W.S. Moffatt | RJA | 1897 | ILL | ♀ flowers | North Carolina / USA | city, cultivated | 35.5665 | -82.5430 | 3 | RJA | ds_non-UDG |
| CM038305 | R.R. Dreisbach 1005 | RSA | 1922 | CM | ♂ flowers | Pennsylvania / USA | waste places | 40.0320 | -75.1313 | 4 | RSA | ds_non-UDG |
| TENN-V-0088565 | E.M. Browne & E.T. Browne Jr. 8949 | RJA | 1964 | TENN | unopened flowers | Kentucky / USA | roadside, highway | 37.3811 | -82.2823 | 3 | RBO | ds_non-UDG |
| AMD.100949 | H. Dennÿs | RSA | 1966 | AMD | unopened flowers | Ireland | mountain range | 53.2539 | -6.36654 | 3 | RJA | ds_half-UDG |
| L 0859517 | P. Hessel, J. Klein, W.V. Rubers 1951 | RJA | 1968 | L | ♀ flowers | Ireland | in village, roadside | 53.7264 | -7.79312 | 3 | RJA | ds_half-UDG |
| 148703 | Druce 91 | RJA | 1906 | OXF | unopened flowers | Wales / UK | village, waste grounds | 51.5766 | -4.10220 | 3 | RJA | ds_half-UDG |
| 148723 | J.E. Lousley | RJA | 1930 | OXF | ♀ flowers | England / UK | city, waste grounds | 51.3937 | -0.15884 | 3 | RJA | ds_half-UDG |
| 148752 | C. Waterfall | RJA | 1931 | OXF | ♀ flowers | England / UK | village, waste grounds | 53.2090 | -2.87042 | 3 | RJA | ds_half-UDG |
| 148225 | E.F. Warburg 776 | RSA | 1947 | OXF | unopened flowers | Scotland / UK | town, near road | 56.8184 | -5.10846 | 4 | RSA | ds_half-UDG |
| B 10 1201968 | B. Wurzell | RBO - 4x | 1987 | B | unopened flowers | England / UK | city | 51.6056 | 0.06820 | 4 | RBO | ds_half-UDG |
| NCY018629 | R.N. Rendatler | RJA | 1853 | NCY | ♀ flowers | France | city, private garden | 48.6938 | 6.17583 | 3 | RJA | ds_half-UDG |
| E00953073 | O.M. Stewart OS/149/93 | RJC | 1993 | E | unopened flowers | Scotland / UK | road | 54.8379 | -4.04754 | 6 | RJC | ds_half-UDG |
| V.2002.024.082 | A.P. Conolly | RJA | 1977 | NMW | ♂ flowers | England / UK | close to river | 54.9746 | -1.81955 | 4 | RBO | ds_half-UDG |
| CGE00070954 | P.D. Sell & J.E. Raven 67/1754 | RJA | 1967 | CGE | unopened flowers | England / UK | roadside hedge | 52.4300 | 0.30122 | 3 | RJA | ds_half-UDG |
| CGE00070956 | R.J. Bunon 1475 | RJA | 1914 | CGE | ♀ flowers | England / UK | city | 50.8357 | -0.78260 | 3 | RJA | ds_half-UDG |
| 0148282* | H.J.M Bowen | RSA | 1955 | OXF | ♂ flowers | England / UK | village, roadside | 51.5999 | -2.20977 | 4 | RSA | ds_half-UDG |
| P05145876 | A. Bouchon 3540 | RSA | 1920 | P | ♂ flowers | France | botanical garden | 44.8452 | -0.56093 | 4 | RBO | ds_half-UDG |
| BR0000024963299 | F. Boon | RJA | 1919 | BR | unopened flowers | Belgium | city | 51.1390 | 4.62976 | 3 | RJA | ds_half-UDG |
| CP0048291 | - | RJA | 1870 | C | ♀ flowers | Denmark | city | 55.7585 | 12.5752 | 3 | RJA | ds_half-UDG |
| L 0807610 | Wanser 248 | RSA | 1910 | L | vegetative | Netherlands | city, road | 52.0713 | 4.28091 | 4 | RSA | ds_half-UDG |
| WAG.1547045 | P. Buwalda 1193 | RJA | 1932 | WAG | ♀ flowers | Netherlands | botanical garden | 53.1805 | 6.60177 | 3 | RJA | ds_half-UDG |
| GJO0088094 | L. Kristof | RJA | 1872 | GJO | one branch with ♂ flowers & one branch with fruits | Austria | botanical garden | 48.1915 | 16.3835 | 3 | RJA | ds_half-UDG |
| 593599 | J. Appl | RJA | 1934 | BRNU | ♀ flowers | Czech Republic | city | 50.6391 | 13.8345 | 3 | RJA | ds_half-UDG |
| V-1025319 | F. Schmidt & P. Von Glehn | RSA | 1860 | UPS | fruits | Russia | village | 47.0468 | 142.052 | 4 | RSA | ds_half-UDG |
| KRAM00033636-V | K. Nowak 211 | RJA | 1988 | KRAM | fruits | Poland | city, roadside | 49.6591 | 19.1973 | 6 | RJC | ds_half-UDG |
| CSFI005348 | L. Xuegen 201824 | RJA | 1958 | CSFI | unopened flowers | Guangdong / China | mountain range, waterside | 24.5214 | 115.582 | 1 | RJA | ds_half-UDG |
| CSFI005352 | J80, 12038 | RJA | 1982 | CSFI | fruits | Zhejiang / China | valley foothills | 30.4920 | 119.614 | 1 | RJA | ds_half-UDG |
| IBK00019687 | T. Peixiang 63853 | RJA | 1959 | IBK | unopened flowers | Hunan / China | mountain range | 26.8859 | 112.611 | 1 | RJA | ds_half-UDG |
| IBK00019738 | G. Xipeng 53467 | RJA | 1933 | IBK | fruits | Guangdong / China | old mine pit | 24.7966 | 113.270 | 1 | RJA | ds_half-UDG |
| IBK00019741 | H.Y. Liang 61207 | RJA | 1931 | IBK | unopened flowers | Guangdong / China | streamside | 23.0224 | 113.768 | NA | RJA | ds_half-UDG |
| IBK00391085 | Lingchuan County Census Team 450323130619015LY | RJA | 2013 | IBK | unopened flowers | Guangxi / China | broadleaf forest, along ditches | 25.1718 | 110.414 | 1 | RJA | ds_half-UDG |
| IMC0039113 | L. Zhengyu 183005 | RJA | 2008 | IMC | unopened flowers | Chongqing / China | mountain streamside | 29.5551 | 106.554 | 1 | RJA | ds_half-UDG |
| PE00235475 | Z. Cao, J. Jin & Hamiti 19850900 | RJA | 1982 | PE | unopened flowers | Hebei / China | city | 39.9368 | 116.415 | 1 | RJA | ds_half-UDG |
| PE00235927 | 236-6 teams 638 | RJA | 1974 | PE | vegetative | Fujian / China | NA | 27.7565 | 118.022 | 1 | RJA | ds_half-UDG |
| PE00242750 | Plant survey team, Henan Academy of Sciences T0614 | RJA | 1986 | PE | fruits | Hubei / China | mountain area, riverbank | 30.8924 | 115.745 | 1 | RJA | ds_half-UDG |
| PE00384031 | S. Lan 483 | RJA | 1985 | PE | unopened flowers | Yunnan / China | city, cultivated | 26.0800 | 103.180 | 1 | RJA | ds_half-UDG |
| E00006538 | G. Forrest 8114 | RJA | 1912 | E | unopened flowers | Yunnan / China | volcanic mountain flank | 25.4653 | 98.3786 | 1 | RJA | ds_half-UDG |
| ILLS241904 | C. Guo 20061-440-3 | RJA | 2006 | ILLS | - | Shandong / China | city | 35.7632 | 117.503 | 1 | RJA | ds_half-UDG |
| P05172070 | K. Yao 9294 | RJA | 1985 | P | ♂ flowers | Jiangxi / China | roadside, on slopes | 27.6525 | 114.593 | 1 | RJA | ds_half-UDG |
| Z-000234655 | Sino American Guizhou Botanical Expedition 929 | RJA | 1986 | Z | ♂ flowers | Guizhou / China | riverbank, roadside, mountain range | 27.8280 | 108.299 | 1 | RJA | ds_half-UDG |
| P05000011 | Y. Tateishi 365 | RJFC | 1971 | P | fruits | Honshu / Japan | mountain area | 35.3865 | 138.748 | 6 | RJC | ds_half-UDG |
| E00849600 | N. Fujii, S. Hashiba, Y. Sakurazawa, K. Watanabe, E. Maki & T. Sugawara 4101658 | RJA | 2004 | E | fruits | Honshu / Japan | river valley, forest margin | 35.8986 | 137.657 | 1 | RJA | ds_half-UDG |
| BS33853 | S. Sudo 3853 | RJC | 1975 | GMNJ | unopened flowers | Honshu / Japan | mountain range | 37.0583 | 139.097 | 6 | RJA | ds_half-UDG |
| FKSE13719 | M. Sato | RJA | 2001 | FKSE | unopened flowers | Honshu / Japan | paddy fields | 37.6471 | 140.628 | 3 | RJA | ds_half-UDG |
| FKSE7330 | H. Sase 78-278 | RJA | 1989 | FKSE | ♀ flowers | Honshu / Japan | city | 34.1040 | 131.293 | 2 | RJA | ds_half-UDG |
| MBK4724 | S. Okumiya 3749 | Rumex madaio | 1988 | MBK | vegetative | Shikoku / Japan | village | 33.8041 | 133.368 | 5 | RSA | ds_half-UDG |
| C1-167001 | T. Kobayashi 31886 | RSA | 1998 | HYO | unopened flowers | Honshu / Japan | mountain range, swamp | 40.6479 | 140.854 | 4 | RSA | ds_half-UDG |
| NY03759254 | A.S. Pease | RJA | 1906 | NY | fruits | Massachusetts / USA | city, waste grounds | 42.3427 | -71.0549 | 3 | RJA | ds_half-UDG |
| ILLS00078672 | H.S. Papon | RSA | 1931 | ILLS | ♂ flowers | Tennessee / USA | mountain range, national park | 35.7898 | -83.4556 | 4 | RSA | ds_half-UDG |
| CM038282 | N. McClintock | RJA | 1931 | CM | fruits | Pennsylvania / USA | city | 40.2478 | -79.2331 | 1 | RJR | ds_half-UDG |
| ILL00129912 | D.S. Seigler 8814 | RJA | 1974 | ILL | unopened flowers | West Virginia / USA | rest area, road | 38.1915 | -81.4904 | 3 | RJA | ds_half-UDG |
| ASU0123287 | S.F. Glassman 8623 | RJA | 1968 | ASU | ♀ flowers | Illinois / USA | city, cultivated | 41.9995 | -87.8748 | 6 | RJC | ds_half-UDG |
| NCU00031364 | M.F Johnson, W. Stepka, J.D. Reynolds 3654 | RJA | 1970 | NCU | ♂ flowers | Virginia / USA | moist roadside | 37.3672 | -77.8694 | 3 | RBO | ds_half-UDG |
| ASU0123283 | M.J. Roberts 4458 | RJA | 1974 | ASU | ♀ flowers | Ohio / USA | along stream | 38.6093 | -82.6261 | 3 | RJA | ds_half-UDG |
| 0084515MOR | D. Ladd 2263 | RJA | 1977 | MOR | vegetative | Vermont / USA | along road | 44.3915 | -72.6488 | 3 | RJA | ds_half-UDG |
| GA140495 | W.T. Gillis 14949 | RJC | 1978 | GA | fruits | Michigan / USA | city, university campus | 42.7132 | -84.4760 | 6 | RJC | ds_half-UDG |
| ASU0123290 | W.L. Halvorson 78-134 | RJA | 1978 | ASU | ♀ flowers | Rhode Island / USA | city, waste places | 41.4842 | -71.5316 | 3 | RJA | ds_half-UDG |
| DOV0030393 | J.R. Grant 89-00503 | RSA | 1989 | DOV | fruits | Maryland / USA | roadside | 39.0015 | -77.0476 | 3 | RBO | ds_half-UDG |
| G00421258 | R. Cayouette 8185 | RJA | 1966 | G | ♀ flowers | Quebec / Canada | city, cemetery | 46.8183 | -71.1665 | 3 | RJA | ds_half-UDG |
| 418826 | M.J. Stufflebeam & D.A. Thompson | RJA | 1971 | CAS | fruits | California / USA | city | 37.7384 | -122.407 | 3 | RJA | ds_half-UDG |
| 03262268 | J.A. & H.F. Ruth | RJA | 1914 | US | ♀ flowers | New Jersey / USA | city | 40.8628 | -74.1620 | 3 | RJA | ds_half-UDG |
| 03262270 | J.A. Niuewland | RJA | 1912 | US | fruits | Indiana / USA | city | 41.7083 | -86.2558 | 3 | RJA | ds_half-UDG |
| MT00290717 | B. Boinier | RSA | 1942 | MT | ♂ flowers | Quebec / Canada | city, garden | 45.5567 | -73.5563 | 4 | RSA | ds_half-UDG |
| CAN10161050 | M.L. Fernald, B. Long, D.H. Linder 21136 | RJA | 1920 | CAN | unopened flowers | Nova Scotia / Canada | roadside, waste ground | 43.8333 | -66.1167 | 3 | RJA | ds_half-UDG |
| LY0005246 | R. Bonaparte | RSA | 1904 | LY | ♂ flowers | North Carolina / USA | city, cultivated | 35.5405 | -82.5523 | 4 | RSA | ds_half-UDG |
| P04912694 | J.A. Drushel | RJA | 1933 | P | vegetative | New York / USA | city | 40.6223 | -74.0769 | 4 | RSA | ds_half-UDG |
| 2129353 | A.N. Steward 7581 | RSA | 1953 | LD | ♂ flowers | Oregon / USA | road along river | 45.4745 | -123.843 | 4 | RSA | ds_half-UDG |
| 03262263 | L.E. James 2705 | RJA | 1955 | US | unopened flowers | Ontario / Canada | city | 42.7703 | -81.1789 | 3 | RJA | ds_half-UDG |
| L14-2022-13 | I. Alexeev & V. Makarov | RJA | 1975 | C | unopened flowers | Russia | city | 52.4184 | 34.7942 | 3 | RBO | ds_half-UDG |
| SRR25018594 | J. Chen | RJA | 2023 | NA | - | Hubei / China | city | 30.2908 | 108.937 | 1 | RJA | modern, no treatment |
| SRR25018595 | J. Chen | RJA | 2023 | NA | - | Shandong / China | NA | 36.0310 | 118.726 | 1 | RJA | modern, no treatment |
| SRR25018596 | J. Chen | RJA | 2023 | NA | - | Hebei / China | city | 39.9123 | 116.394 | 1 | RJA | modern, no treatment |

**Table S2**. Unique dual indices used to discriminate between different libraries. All libraries in a batch were equimolarly pooled and sequenced in parallel in a single run on an Illumina NovaSeq platform at Genewiz (Azenta Life Sciences, Leipzig, Germany) using a 2x150bp pair-end kit.

| **Library name** | **i7 Index Sequence** | **i5 Index Sequence** | **Library Batch** |
| --- | --- | --- | --- |
| FI58969 | TGGTTGAC | TAACTATC | L1 |
| B100481650 | TGGTATGA | CTATTACC | L1 |
| ZU6 | GTTAGCGG | TGATAGGT | L1 |
| BEOU46519 | TATATGCC | TCGATCCT | L1 |
| P500007 | TAACGTCG | CAACCGAT | L1 |
| LY0005293 | TAATCTGC | GGTCAGCA | L1 |
| ZG60886 | TGGTCCTG | TGGATAGC | L1 |
| TUR567401 | ATAAGACT | TGAGAGCA | L1 |
| B101201970 | AACCTGCC | CTGGTAGG | L1 |
| BEOU37718 | TTGGAGCT | AGTTGGTT | L1 |
| BR12293353 | GTACTCGA | CCGGTATT | L1 |
| B101201981 | CTTCGTTA | CCTTACTG | L1 |
| GJO88103 | TGGCCGAG | GCGATCAA | L1 |
| L0768025 | GGCGTTGC | AACCGAAC | L1 |
| P04961759 | ATTGAAGG | GCGACGAC | L1 |
| BEOU27311 | GCTTGATG | AACCATGC | L1 |
| M0323153 | CGGACGGC | CAACCAGG | L1 |
| B101201971 | TTCGATGA | CCAATCCA | L1 |
| V174556 | CGCATCAT | TTAGGAGG | L1 |
| CGE70939 | ATGAATCT | GGAACTGG | L1 |
| MA782918 | GGTTAGTC | TGATGAGA | L1 |
| G00421221 | CCAAGTCA | TGCGGACC | L1 |
| FLAS8116 | GTCTTGGC | GGACGCCT | L2 |
| FLAS31625 | GGCGGTAT | GTTGCTGG | L2 |
| TENN0088557 | GTCGCTAG | ACCAGACC | L2 |
| ILL00129901 | CCTGGTAC | AACGCATC | L2 |
| MIN379045 | GGTTCAGC | CTAACTTA | L2 |
| ILL0129908 | CGTATTGC | ACCATTAC | L2 |
| RSA0503144 | CCGTACTA | GATACTTG | L2 |
| RSA0503137 | TTACTTCT | TTCGTTGA | L2 |
| 0084518MOR | TGAAGGTT | GTTGCCAA | L2 |
| US3262258 | AACCAACG | GAACCATT | L2 |
| UT0073333 | GCATTGGT | GTATTCGG | L2 |
| 0084512MOR | TGGTAATT | CCGAACGG | L2 |
| CM37781 | CCGTCTGC | TGGTAGTC | L2 |
| RSA0503141 | AGCCAATC | CGAAGTAG | L2 |
| KANU351723 | ATGCCGGA | CCGGTTAG | L2 |
| GA140493 | GCAGTCCA | TCGGTCTC | L2 |
| UT0073288 | TCATTCCG | CAGAACTT | L2 |
| TENN0087579 | GCGGTTCT | GGTCAGCA | L2 |
| NY3759204 | TGGTTGAC | CGAGAGTT | L2 |
| TENN0088577 | TGGAGTAC | ACCAATCA | L2 |
| ILLS00078641 | CCAAGTCA | GGTTGCAT | L2 |
| KANU00082404 | GTTCAGGC | GTTGCTGG | L2 |
| B101201978 | TTGGCAGA | TTCTCGTT | L3 |
| B101201967 | GCGTTAGC | AGAAGAGC | L3 |
| MBK292730 | TGGTAATT | AATGAGAT | L3 |
| TOYA42355 | TCTATTCG | TGCTCATG | L3 |
| B101201963 | CCGTACTA | AACCATGC | L3 |
| TOYA86770 | TTGACTAG | TCGGTCTC | L3 |
| B101201980 | CGCTCGAC | GTTGGAAC | L3 |
| TOYA107657 | GAACGCTG | AGGTCGCG | L3 |
| C1080297 | AGAATAAG | ACCAGACC | L3 |
| P0500037 | TTGGAGCT | CCTTGGAA | L3 |
| TOYA86153 | CGGACGGC | GCGACGAC | L3 |
| B100426659 | GTACTCGA | AACGAAGT | L3 |
| B101201982 | GCTGGTCA | TAACTATC | L3 |
| TOYA84947 | GAAGTCGT | CCGCTACG | L3 |
| FKSE69323 | GGCGTTGC | TTAGATTC | L3 |
| ZU7 | GGTAGTTG | TCTTGCCT | L3 |
| TOYA83438 | ATGCCGGA | GATACTTG | L3 |
| FKSE7311 | TAACGTCG | GCTTCTTA | L3 |
| WU0131126 | CGCATCAT | GGAACTGG | L3 |
| TOYA55320 | GGCGGTAT | ATAGTCAG | L3 |
| GONA | GAGAGGCC | GTTGCCAA | L3 |
| CM297915 | TGACTCAA | ATTGGTAA | L3 |
| LY0005301 | ACTTAGTA | TCTTGCCT | L0 |
| TENN0088565 | AACCTAGA | TGGAGATG | L0 |
| 128371 | AACCTCAG | AAGAATGG | L0 |
| P04940026 | AACTACCT | GCTTCTTA | L0 |
| CM435363 | AACCGAAC | AAGAAGAC | L0 |
| M0323149 | AACGAAGT | AACGACTA | L0 |
| P04555472 | AAGAACCA | CCGAGACT | L0 |
| 0084521MOR | ACCAGACC | AGGCTTCT | L0 |
| MA509322 | CAATCCGG | CTCTGCTG | L0 |
| NY3759272 | AATCCATG | GGCTTGCT | L0 |
| KANU00324585 | ACGAACTT | GGTCGTCC | L0 |
| BEOU55742 | AAGAGAAT | AACCGAAC | L0 |
| DOV30381 | CAACCATC | TGCGGACC | L0 |
| KANU363875 | AACCTAGA | AACCTCAG | L0 |
| MIN103352 | ATTGAAGG | GTTCTATA | L0 |
| FI065948 | AAGAACCA | TGAGAGCA | L0 |
| ILL00129906 | ACTTAGTA | GTCGCGCT | L0 |
| B101201979 | AACGACTA | TGACGTTC | L0 |
| CM38305 | ACCAGACC | AAGAACCA | L0 |
| TAA0091032 | AACCGAAC | TCGATCCT | L0 |
| AMD100949 | TGGACGCA | AACCGAAC | L4 |
| U1530588 | TCATTCCG | GGACGCCT | L4 |
| 00148703N | TAATCTGC | TTCTCGTT | L4 |
| 00148723P | CCGTCTGC | AAGAATGG | L4 |
| 00148752R | CCTCGACG | TGATAGGT | L4 |
| 00148225M | TTGGCAGA | TTGACGCA | L4 |
| B101201968 | AACCTAGA | GTTGGAAC | L4 |
| NCY018629 | AACCAACG | GGCTTGCT | L4 |
| E00953073 | CCTGCCAA | GTTGCTGG | L4 |
| V2002024082 | TGGCCGAG | GCTTCTTA | L4 |
| CGE00070954 | CCATACCT | CTAACTTA | L4 |
| CGE00070956 | CCGTACTA | AACCAACG | L4 |
| OXF0148282P | CGTTCGCT | ACCATTAC | L4 |
| P05145876 | ACCAAGAT | CCTAGCAT | L4 |
| BR24963299 | AGAACCGG | TTAGATTC | L4 |
| CP0048291 | AAGAGAAT | CAGCCGTA | L4 |
| L3190307 | ATAAGACT | TGCGGACC | L4 |
| WAG1547045 | TAACGTCG | GTTGCCAA | L4 |
| GJO0088094 | TTGGAGCT | AATGAGAT | L4 |
| 593599 | TGGTAATT | AACCAGAA | L4 |
| V-1025319 | ACGAACTT | GAACCATT | L4 |
| KRAM00033636-V | AACGACTA | TAGTCTAC | L4 |
| CSFI005348 | TGGCCGAG | ACTCTTGG | L5 |
| CSFI005352 | GTATCTGA | AAGAATGG | L5 |
| IBK00019687 | CCGTCTGC | TATGCGCC | L5 |
| IBK00019738 | CCGCCATT | CCTTACTG | L5 |
| IBK00019741 | TATATGCC | CGAAGTAG | L5 |
| IBK00391085 | CCTCGACG | CAACCGAT | L5 |
| IMC0039113 | GCGCGGTA | CGTTGGCG | L5 |
| PE00235475 | TTCGATGA | TCCATCGC | L5 |
| PE00235927 | CAAGCTAA | ACCTTCGG | L5 |
| PE00242750 | CCTGACGG | GGTAGCCA | L5 |
| PE00384031 | CCGCCATT | CAACCGAT | L5 |
| E00006538 | AACCGAAC | TTCTACTC | L5 |
| ILLS241904 | AAGAACCA | ACCAGACC | L5 |
| P05172070 | AACGAAGT | GAACGTAC | L5 |
| Zu10 | GTCGCTAG | GTTGAATT | L5 |
| FKSE7330 | TTCAGCAG | GTATTCGG | L5 |
| MBK4724 | GGTTAGTC | CTGGTAGG | L5 |
| C167001 | AACCGGTT | ACTCTTGG | L5 |
| P05000011 | GCTTGATG | CGAAGTAG | L5 |
| E00849600 | ACCAGACC | AACCATGC | L5 |
| BS0033853 | TGGAGTAC | AACCGCCA | L5 |
| FKSE13719 | CAATCCGG | GTCGATAA | L5 |
| OXF00148282a | ATTGAAGG | AACGAAGT | L6 / technical replicate |
| 0084518MORa | AGAGAACG | TAACTATC | L6 / technical replicate |
| RSA0503141a | GAAGTCGT | AACGACTA | L6 / technical replicate |
| ILL00129906a | GCGTTAGC | GGCTATGC | L6 / technical replicate |
| BEOU55742a | GAACGCTG | CCGGTATT | L6 / technical replicate |
| BEOU46519a | CGCATCAT | ATAGTCAG | L6 / technical replicate |
| P04940026a | TGGTCCTG | CCTTACTG | L6 / technical replicate |
| GONAa | CAAGCTAA | GCTGCGGA | L6 / technical replicate |
| G00421221a | TGGTATGA | ACCTTCGG | L6 / technical replicate |
| P0500037a | TTGGCTCC | GTATCTGA | L6 / technical replicate |
| Zu10a | GTTCAGGC | ACCAATCA | L6 / technical replicate |
| OXF00148282b | GGCGTTGC | GGTAGCCA | L6 / technical replicate |
| 0084518MORb | CGCTCGAC | TGGAGATG | L6 / technical replicate |
| RSA050314b | CGGACGGC | TGGATAGC | L6 / technical replicate |
| ILL00129906b | GTATCTGA | CCGGTTAG | L6 / technical replicate |
| BEOU55742b | TGGCCGAG | AACCAACG | L6 / technical replicate |
| BEOU46519b | TTCAGCAG | AACCAGAA | L6 / technical replicate |
| P04940026b | ATGAATCT | GGTAGCCA | L6 / technical replicate |
| GONAb | CTTCGTTA | AGAAGAGC | L6 / technical replicate |
| G00421221b | AACCAGAA | TGGTAGTC | L6 / technical replicate |
| P0500037b | AACGACTA | AATGAGAT | L6 / technical replicate |
| Zu10b | GTTAGCGG | CCTCGATA | L6 / technical replicate |
| NY3759254 | GGTTCAGC | GGTCGTCC | L7 |
| ILLS00078672 | TGGACGCA | CGAAGTAG | L7 |
| CM38282 | AGCCAATC | GTCGCGCT | L7 |
| ILL00129912 | TGGCGTTA | AACCATGC | L7 |
| ASU123287 | GTTCAGGC | AACGACTA | L7 |
| NCU31364 | CCTGCCAA | TGACGTTC | L7 |
| ASU0123283 | GCGGTTCT | AACCGCCA | L7 |
| 0084515MOR | GCGCGGTA | CAGCCGTA | L7 |
| GA140495 | ATGCCGGA | TGAGAGCA | L7 |
| ASU0123290 | GTTGCAAC | AAGAATGG | L7 |
| DOV0030393 | CATGGAAG | TAGTCTAC | L7 |
| G00421258 | ACTTAGTA | GGTCAGCA | L7 |
| 418826 | TAATCTGC | AAGAAGAC | L7 |
| 3262268 | GGCGGTAT | GCGACGAC | L7 |
| 3262270 | TGGTCCTG | ATTGGTAA | L7 |
| MT00290717 | CAAGCTAA | TTAGATTC | L7 |
| CAN10161050 | GCATTGGT | TGCGGACC | L7 |
| LY0005246 | TTGGAGCT | TTGCCTGG | L7 |
| P04912694 | GGCGTTGC | AACCTCAG | L7 |
| 2129353 | TTCTGAAC | CAACCAGG | L7 |
| 3262263 | CAATCCGG | TTCTACTC | L7 |
| L14-2022-13 | GTTGACCG | GAAGAGGT | L7 |

**Table S3**. Total allele counts and Jaccard similarity matrix for *R. japonica* across its introduced (Europe, North America) and native regions (China, Japan).

| **Region** | **Total Alleles** | **Europe Jaccard** | **North America Jaccard** | **Japan Jaccard** | **China Jaccard** |
| --- | --- | --- | --- | --- | --- |
| Europe | 2744 | 1 | 0.762 | 0.603 | 0.308 |
| North America | 2620 | 0.762 | 1 | 0.654 | 0.325 |
| Japan | 3063 | 0.603 | 0.654 | 1 | 0.287 |
| China | 3647 | 0.308 | 0.325 | 0.287 | 1 |

**Table S4.** Total allele counts and Jaccard similarity matrix for *R. sachalinensis* across its introduced (Europe, North America) and native regions (Japan).

| **Region** | **Total Alleles** | **Europe Jaccard** | **North America Jaccard** | **Japan Jaccard** |
| --- | --- | --- | --- | --- |
| Europe | 2378 | 1 | 0.920 | 0.858 |
| North America | 2358 | 0.920 | 1 | 0.864 |
| Japan | 2431 | 0.858 | 0.864 | 1 |

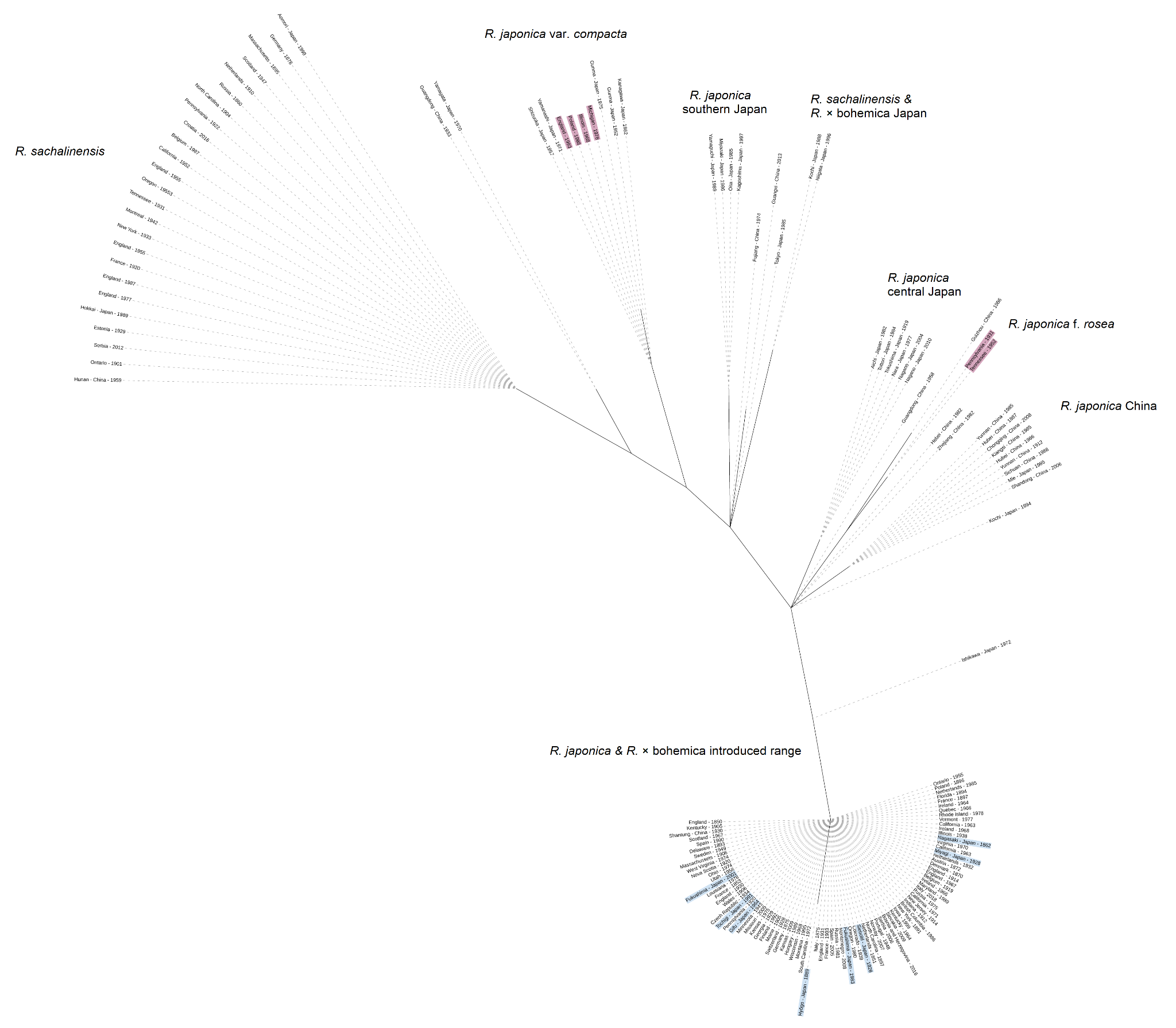

**Figure S1.** Maximum likelihood phylogenetic tree of 153 Japanese knotweed *s*.*l*. accessions constructed using 27 chloroplast SNPs.

Labels highlighted in blue denote *R. japonica* specimens from Japan that represent the putative native source of the invasion. Labels highlighted in pink denote individuals of *R. japonica* var. *compacta* and R. *japonica* f. *rosea* from Europe and North America.

Supplementary text for Figure S1

The phylogeny based on only 27 plastid SNPs showed a structure broadly consistent with the topology recovered from the full SNP dataset (Fig. S1). All *R. japonica* individuals from Europe and North America clustered within a restricted portion of the tree and shared a single, closely related plastid type, indicating a strong maternal founder effect. *Reynoutria* × *bohemica* from Europe and North America were placed within the same plastid clade, supporting a shared maternal origin and suggesting that *R. japonica* most likely served as the maternal parent during hybrid formation. All *R. sachalinensis* individuals with the exception of one specimen from southwestern Japan, formed a distinct plastid clade clearly separated from the invasive *R. japonica* lineage. The three *R.* × *bohemica* hybrids from the UK and France clustered within the *R. sachalinensis* plastid clade and shared an identical plastid type, indicating that *R. sachalinensis* served as the maternal parent in these hybridization events. *Reynoutria japonica* var. *compacta* accessions from both native and introduced regions formed a single plastid clade distinct from both the *R. sachalinensis* lineage and the invasive *R. japonica* plastid clade, supporting its status as a genetically differentiated maternal lineage within Japan. Interestingly, the *R.* × *bohemica* specimen from Japan and the *R. sachalinensis* specimen from southwestern Japan shared the same plastid type, which traces back to a maternal lineage also found in *R. japonica* from southern Japan and two Chinese *R*. *japonica* accessions. Several plastid types were detected for *R*. *japonica* in its native range. The *R. japonica* f. *rosea* plastid traced back to a common maternal ancestor shared with *R. japonica* from southwest China (Fig. S1).
